# Progressive and accurate assembly of multi-domain protein structures from cryo-EM density maps

**DOI:** 10.1101/2020.10.15.340455

**Authors:** Xiaogen Zhou, Yang Li, Chengxin Zhang, Wei Zheng, Guijun Zhang, Yang Zhang

**Author notes:** Correspondence should be addressed to Y. Z.

## Abstract

Progress in cryo-electron microscopy (cryo-EM) has provided the potential for large-size protein structure determination. However, the solution rate for multi-domain proteins remains low due to the difficulty in modeling inter-domain orientations. We developed DEMO-EM, an automatic method to assemble multi-domain structures from cryo-EM maps through a progressive structural refinement procedure combining rigid-body domain fitting and flexible assembly simulations with deep neural network inter-domain distance profiles. The method was tested on a large-scale benchmark set of proteins containing up to twelve continuous and discontinuous domains with medium-to-low-resolution density maps, where DEMO-EM produced models with correct inter-domain orientations (TM-score >0.5) for 98% of cases and significantly outperformed the state-of-the-art methods. DEMO-EM was applied to SARS-Cov-2 coronavirus genome and generated models with average TM-score/RMSD of 0.97/1.4Å to the deposited structures. These results demonstrated an efficient pipeline that enables automated and reliable large-scale multi-domain protein structure modeling with atomic-level accuracy from cryo-EM maps.

## INTRODUCTION

Single-particle cryo-electron microscopy (cryo-EM) has emerged as a powerful means in modeling macromolecular structures at near-atomic resolution^1,2^. High resolution density maps allow direct construction of atomic structures with limited conformation sampling using software programs traditionally used for X-ray crystallography^3-5^. However, the performance of the programs is poor when the resolution of density map is relatively low (e.g., >3Å)^5,6^. For these challenging cases, a common approach is to fit a homologous structure into the density map, followed by atom-level structural refinement^7-9^. However, the success of the approach highly depends on the quality of the starting models, while many proteins have no previously solved structures of homologous proteins.

The difficulty is particularly crucial for multi-domain proteins consisting of multiple structurally autonomous subunits. In fact, multi-domain proteins are common in nature and statistics has shown that more than two-thirds of prokaryote proteins and four-fifths of eukaryote proteins are composed of two or more domains^10^; but only one third of structures in the PDB contain multiple domains (**Supplementary Fig. 1a**). Due to the lack of multi-domain templates and the difficulty for *ab initio* domain orientation modeling, the field of computational structural biology has traditionally focused on the study of individual domains, including the community-wide CASP experiments which assess the quality of protein structure predictions mainly on the individual domains^11^. Therefore, although cryo-EM provides a strong potential for determining large-size proteins^2^ and there are considerably more multi-domain proteins in the Electron Microscopy Data Bank (EMDB)^12^ than that in the PDB (**Supplementary Fig. 1b**), it is usually difficult to use homology modeling to create appropriate frameworks for the density-map fitting and structural refinements of multi-domain proteins. These factors impose a significant challenge for cryo-EM based multi-domain structure modeling and are probably the important reasons that only less than half of the cryo-EM density map in the EMDB have atomic structures (**Supplementary Fig. 1c**). An additional barrier in large-scale cryo-EM structural modeling is that almost all structure fitting and refinement tools are not fully automated even with given homologous models. For example, many approaches require human-interventions in initial model-to-map fitting, a procedure that often significantly impacts quality of the final models^13^. Hence, developing advanced cryo-EM methods that can automatically and yet reliably assemble multi-domain structures becomes increasingly urgent given the rapid progress of cryo-EM structural biology.

In this study, we proposed a novel automated approach (termed DEMO-EM, **Fig. 1**) to create accurate complex structure models for multi-domain proteins from cryo-EM density maps. The pipeline can start from either experimentally determined domain structures or amino acid sequences, where in the latter case the domain split and individual domain structure modeling are performed with FUpred^14^ and I-TASSER^15^, respectively. To systematically examine the strength and weakness, DEMO-EM was tested on a large-scale benchmark dataset consisting of various numbers of continuous and discontinuous domains over synthesized and experimental density maps. The results demonstrate significant advantages of DEMO-EM for cryo-EM guided domain structure assembly and refinement compared to the start-of-the-art approaches in the field. The source codes and datasets of DEMO-EM programs, together with an online server, are publicly available at https://zhanglab.ccmb.med.umich.edu/DEMO-EM/.

**Figure 1.**
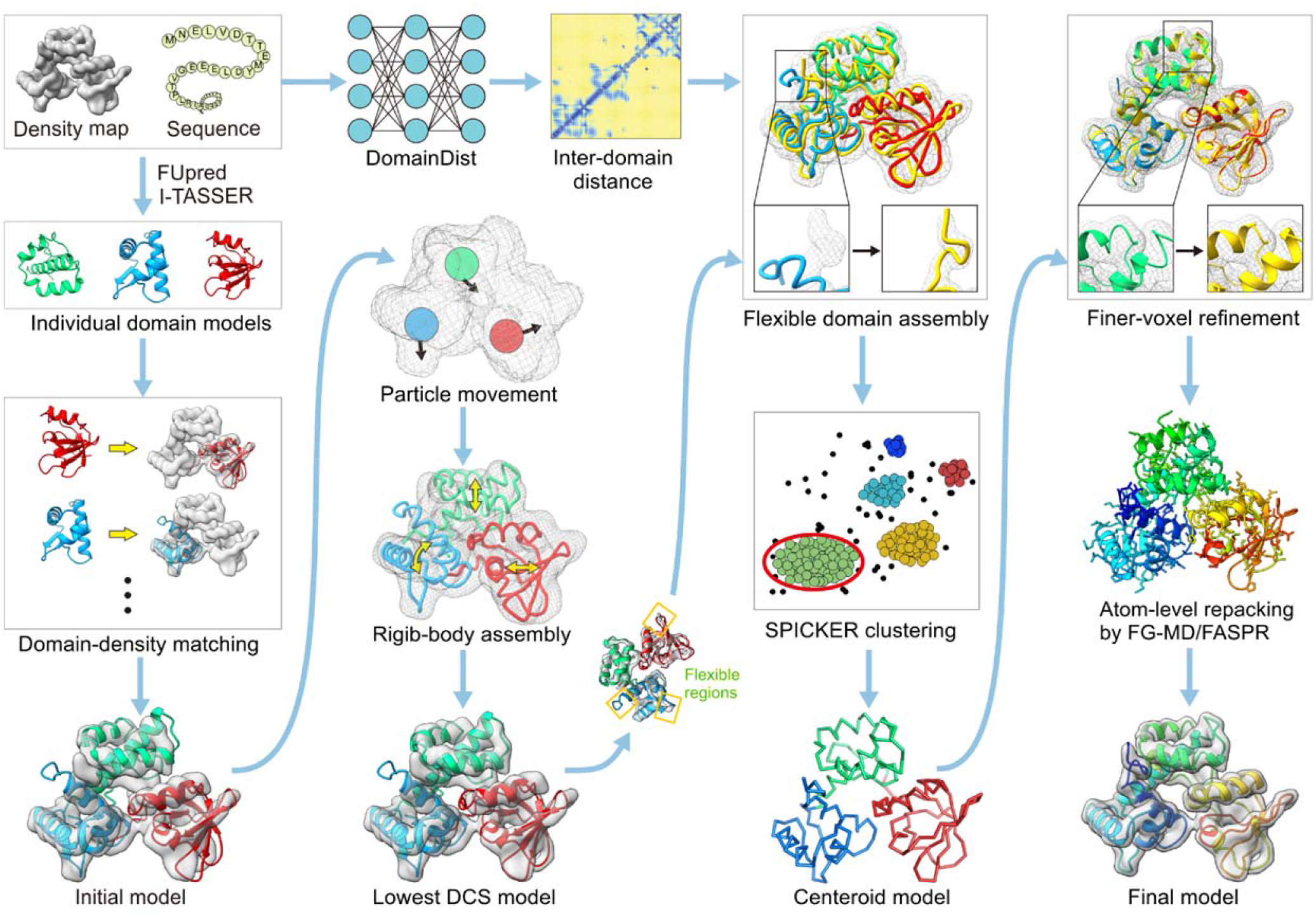
Flowchart of DEMO-EM illustrated with a 3-domain protein from the *iron-dependent regulator of mycobacterium tuberculosis* (PDBID: 1fx7A). Starting from the query sequence, domain boundaries and models of each domain are first predicted by FUpred^14^ and I-TASSER^15^, respectively. Meanwhile, inter-domain distances are predicted with a deep convolutional neural-network predictor DomainDist. Second, each of the domain models is independently fit into the density map by quasi-Newton searching. Third, the initial full-length models are optimized by a two-step rigid-body REMC simulation to minimize the DCS between the density map and full-length model (Eq. 1). Fourth, the lowest DCS model selected from the rigid-body assembly simulations undergoes flexible assembly with atom-, segment-, and domain-level refinements using REMC simulation guided by the DCS and inter-domain distance profiles coupled with a knowledge-based force field, with the resulting decoy conformations clustered by SPICKER^17^ to obtain a centroid model. Finally, the flexible assembly simulation is performed again for the full-atomic model with constraints from centroid models adding to the energy, and the final model is created from the lowest energy model after side-chain repacking with FASPR^18^ and FG-MD^19^.

## RESULTS

### Multi-domain structure construction from synthesized density maps

#### Benchmark data setting

To evaluate DEMO-EM, we collected a set of 357 non-homologous proteins from the PDB with the domain boundary assigned by DomainParser^20^ (or SCOPe^21^ and CATH^22^ when available). The length of these proteins ranges from 99 to 1,693 residues with 2-12 domains. The density maps of these proteins are simulated according to the experimental structures by EMAN2^23^, with the resolution randomly selected from 2 to 10 Å and a grid spacing of 1 Å/voxel (**Supplementary Fig. 2a**). Two separate tests are performed to assemble experimental domain structures extracted from the full-length target structures and domain models predicted by I-TASSER^15^. For experimental domain structure assembly, all domains were randomly rotated and translated as rigid bodies before assembly. When using I-TASSER to model the domain structures, all homologous templates of a sequence identity >30% to the query have been excluded; this resulted in domain models with variable quality and the TM-score ranging from 0.19 to 0.97 (see **Supplementary Fig. 2b** for a histogram of TM-score distribution). Since we focus mainly on examining the domain assembly ability of DEMO-EM, we excluded the proteins with any domain of TM-score^24^ below 0.5 in the second benchmark test, to eliminate the negative impact from incorrect domain models. This resulted in 229 proteins and the average TM-score for them is 0.76.

#### Overall results of DEMO-EM modeling

**Table 1** and **Figure 2** present a summary of the DEMO-EM models on both benchmark tests. When the experimentally determined domain structures are used, DEMO-EM was able to assemble nearly perfect full-length models for almost all the targets, which resulted in an average TM-score=0.99 and RMSD=0.6 Å (**Figs. 2a and 2b**). Importantly, the individual domain structures were well-folded in final full-length models with an average TM-score=0.98 and average RMSD=0.6 Å, despite the fact that atomic structure of full-length models are kept completely flexible in the domain assembly simulations; suggesting that the combination of the inherent DEMO-EM force field and the density-map data is capable to recognize and maintain correct folded domain structures.

**Table 1.** Summary of cryo-EM based domain structure assembly results by different methods.

| Method | 357 proteins with synthesized maps |  |  |  |  |  | 51 proteins with experimental maps |  |  |  |
| --- | --- | --- | --- | --- | --- | --- | --- | --- | --- | --- |
|  | Experimental domain |  | I-TASSER domain |  |  |  |  |  |  |  |
|  | TM-score | RMSD | TM-score | RMSD | TM-score (domain)<sup>a</sup> | RMSD (domain)<sup>b</sup> | TM-score | RMSD | TM-score (domain) | RMSD (domain) |
| MDFF | 0.83 | 7.7 | 0.55 | 17.2 | 0.65 | 6.2 | 0.46 | 22.7 | 0.52 | 9.0 |
| Rosetta | 0.79 | 8.1 | 0.53 | 17.5 | 0.57 | 8.1 | 0.40 | 25.6 | 0.44 | 12.9 |
| DEMO-EM | **0.99** | **0.6** | **0.85** | **6.0** | **0.83** | **4.1** | **0.81** | **7.3** | **0.77** | **4.4** |

<sup>a</sup> TM-score of individual domain models in full-length models. <sup>b</sup> RMSD of individual domains models in full-length models. Bold font highlights the best result in each category.

**Figure 2.**
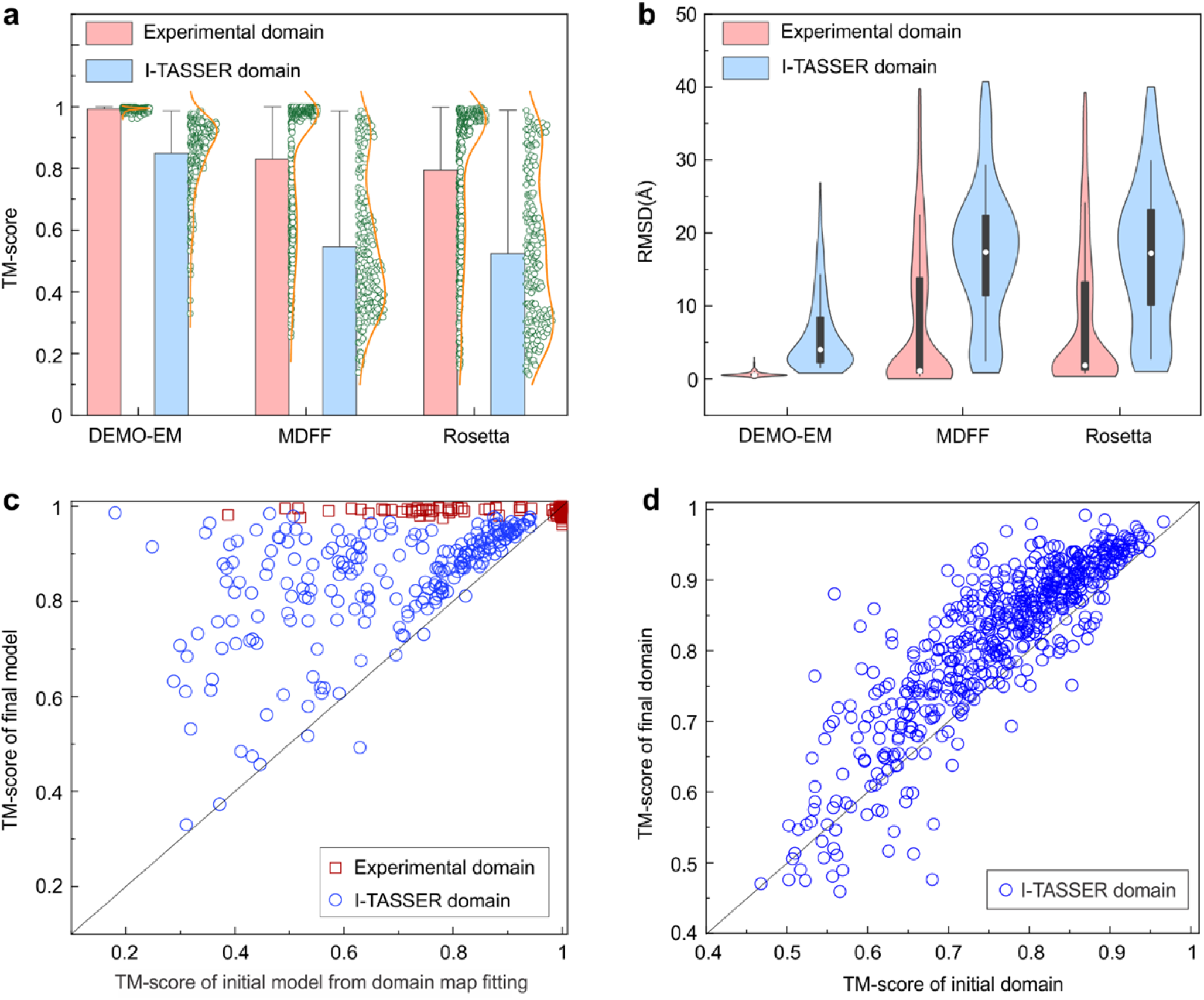
Summary of full-length structural models constructed by different approaches on 357 multi-domain proteins using synthesized density maps. (**a**) Mean and distribution of TM-score for models by DEMO-EM, MDFF and Rosetta, respectively. (**b**) Boxplot and distribution for RMSD of models by DEMO-EM, MDFF, and Rosetta, respectively. (**c**) Head-to-head comparison between TM-score of initial models by domain matching and that of final models after rigid-body assembly, flexible assembly and refinement. (**d**) Comparison between TM-score of individual domain models by I-TASSER and that in final full-length models by DEMO-EM.

When predicted domain models are used, the local structure errors from I-TASSER model can negatively impact the full-length model assembly simulations. Nevertheless, DEMO-EM successfully assembles full-length models with a correct global fold (i.e., TM-score>0.5)^25^ for 97.4% of the test cases (blue histogram in **Fig 2a**). In **Fig 2c**, we presented a head-to-head TM-score comparison between the initial model obtained by matching I-TASSER domains with the cryo-EM maps and the final DEMO-EM model, where the TM-score of the final model was improved in nearly all test cases (with an average TM-score increase from 0.68 to 0.85, corresponding to p-value=6.1E-34 in Student’s t-test). Because creating a high TM-score model requests for correct modeling of both individual domains and inter-domain orientations, the data in **Fig 2c** indicates that DEMO-EM domain assembly simulations could significantly improve the inter-domain orientations. Since the domain structures were kept flexible in DEMO-EM, part of the full-length model TM-score increase may also come from the quality improvement of individual domain structures. To examine this, we presented in **Fig 2d** the TM-score comparison of individual domains by I-TASSER and DEMO-EM, where the TM-score by DEMO-EM was improved in 516 out of the 571 individual domains. On average, the TM-score of individual domains increased from 0.76 to 0.83 with a p-value=8.5E-16 in Student’s t-test, indicating the domain-level structural improvements brought by DEMO-EM are statistically significant.

Interestingly, although the degrees of freedom and searching space usually increases with the number of domains for both domain assembly and full-length model refinement, the quality of the final models by DEMO-EM does not significantly decrease with an increasing number of domains for both experimental and I-TASSER domain model assemblies (**Supplementary Table 1 and 2**). In particular, the full-length models assembled using I-TASSER domain structures obtain an average TM-score of 0.84 for proteins with 3 or more domains, 97% of which have a final model TM-score >0.5. This data demonstrates the ability of DEMO-EM for handling large complex structures of multi-domain proteins. In addition, proteins with discontinuous domains are usually difficult to model as they have several parts separated in sequence which increases the difficulty in individual domain modeling and inter-domain distances prediction. However, DEMO-EM correctly assembled nearly all the cases of discontinuous domains using experimental domain models, with an average TM-score=0.99 identical to that of the continuous domain model assembly (**Supplementary Table 1**). This is probably due to the constraints brought by the additional linkers which help guide the discontinuous domain structure assembly (see **Eq. 4** in Methods and **Supplementary Fig. 13**). When assembling I-TASSER domain models, however, DEMO-EM achieves an average TM-score of 0.83 slightly lower than the continuous domain assembly results (TM-score=0.87, **Supplementary Table 2**), due to the modeling errors of I-TASSER models in the tail and linker regions which affect the packing the segment structures.

#### Control of DEMO-EM with other cryo-EM modeling methods

As a control, we listed in **Table 1** the model results by two widely used methods of MDFF^8^ and Rosetta^13^ for cryo-EM density map guided modeling (**Supplementary Text 1**). Since both methods need to start from full-length models, we built the initial full-length models by fitting each domain model into density maps using Situs^26^, one of the best publicly available structure-density map program. As shown in **Table 1, Figs. 2a** and **2b**, and **Supplementary Figs. 3a and 3b**, DEMO-EM outperformed both control methods by a margin, with the average TM-score of the full-length models with experimental domains 19.3% and 25.3% higher than that of MDFF and Rosetta, respectively. The p-value in student’s t-test is 4.2E-38 and 4.5E-44, respectively, suggesting that the difference is statistically significantly.

When I-TASSER domains were used, the TM-score improvement of DEMO-EM increases to 54.5% relative to MDFF and 60.4% to Rosetta, which corresponds to a student’s t-test p-value of 1.6E-54 and 3.5E-44, respectively. DEMO-EM also made more significant improvement in the domain-level structures. When starting from the I-TASSER domains, DEMO-EM improves the TM-score of individual domains in 89.0% of cases (**Fig. 2d**), while MDFF and Rosetta did so only in 24.0% and 22.1% of the cases respectively (**Supplementary Figs. 3c and 3d**).

#### Why does DEMO-EM outperform control methods?

There are several reasons for the superior performance of DEMO-EM over the control methods. First, the quick quasi-Newton searching process in combination with a space enumeration algorithm as taken by DEMO-EM (see Methods) can correctly match individual domains into the density map and thus generate optimal initial full-length models for the majority of proteins. As shown in **Supplementary Fig. 4a**, the average TM-score of initial full-length models constructed by the quasi-Newton search were 0.97 and 0.69 (where 99.2% and 80.3% had a TM-score >0.5) when starting with experimental and I-TASSER domains, respectively, which are 14.1% and 21.0% higher than those by the start-of-the-art structure-map docking program Situs.

Second, DEMO-EM took a hierarchical process of rigid-body and flexible model assembly simulations to progressively refine the multi-domain structures. In particular, the rigid-body domain assembly process can quickly adjust domain poses based on density maps. When starting with the I-TASSER domains, for example, the average TM-scores of full-length models were improved from 0.69 to 0.78 where the number of cases with TM-score >0.5 increases from 80.0% to 95.2% after the rigid-body assembly step (**Supplementary Fig. 4b**). This domain model improvement helps the subsequent DEMO-EM steps to detect unreasonably folded regions in initial full-length models for more efficient atomic-level structural assembly and refinement simulations.

The last important advantage of DEMO-EM comes from the flexible assembly and refinement stage, which showed a superior ability to improve full-length models and individual domain models simultaneously with the assistance of deep-learning based inter-domain distances predictions when coupled with density-map correlation and inherent DEMO-EM force field. To examine the efficiency of the flexible assembly and refinement process, we feed the same full-length models assembled by the DEMO-EM rigid-body assembly step on the I-TASSER domains into MDFF and Rosetta. Although the better starting model quality resulted in a considerable improved final model for both MDFF and Rosetta, which have the average TM-score of 0.82 and 0.81 respectively, the overall quality is still worse than that of DEMO-EM with an average TM-score 0.85 (**Supplementary Table 3**). Compared to the rigid-body assembled models, the last stage of DEMO-EM simulations improved the TM-score by 9.0%, which is 75% higher than that by MDFF (5.1%) and 133% higher than that by Rosetta (3.8%), demonstrating the efficiency of the atomic-level domain structure refinement of DEMO-EM even starting from the same full-length models.

#### Case studies

In **Figure 3**, we present two illustrative examples showing the construction process of DEMO-EM. For *human xanthine oxidoreductase mutant F3* (PDBID: 2e1qC), a complex protein with 8 continuous domains and 2 discontinuous domains (one of them has three discontinuous segments) using a simulated density map with a medium resolution of 5.3 Å (**Fig. 3a**), one of the domains was initially docked into an incorrect region in the quasi-Newton based search, resulting in an suboptimal full-length TM-score of 0.89 and RMSD=5.4 Å. After the second step of rigid-body assembly, the domain was moved to the correct map space but still with some regions stretched outside the maps in several domains and have the model with a TM-score to 0.95 and RMSD=3.8 Å. At the last step, the flexible simulations refined the overall quality including drawing the exposed loops into the density map, which resulted in a further improved model with TM-score=0.98 and RMSD=2.7 Å. In this case, model quality of the 10 individual domains is also improved with average TM-score/RMSD improved from 0.84/3.2Å to 0.95/1.6Å, respectively.

**Figure 3.**
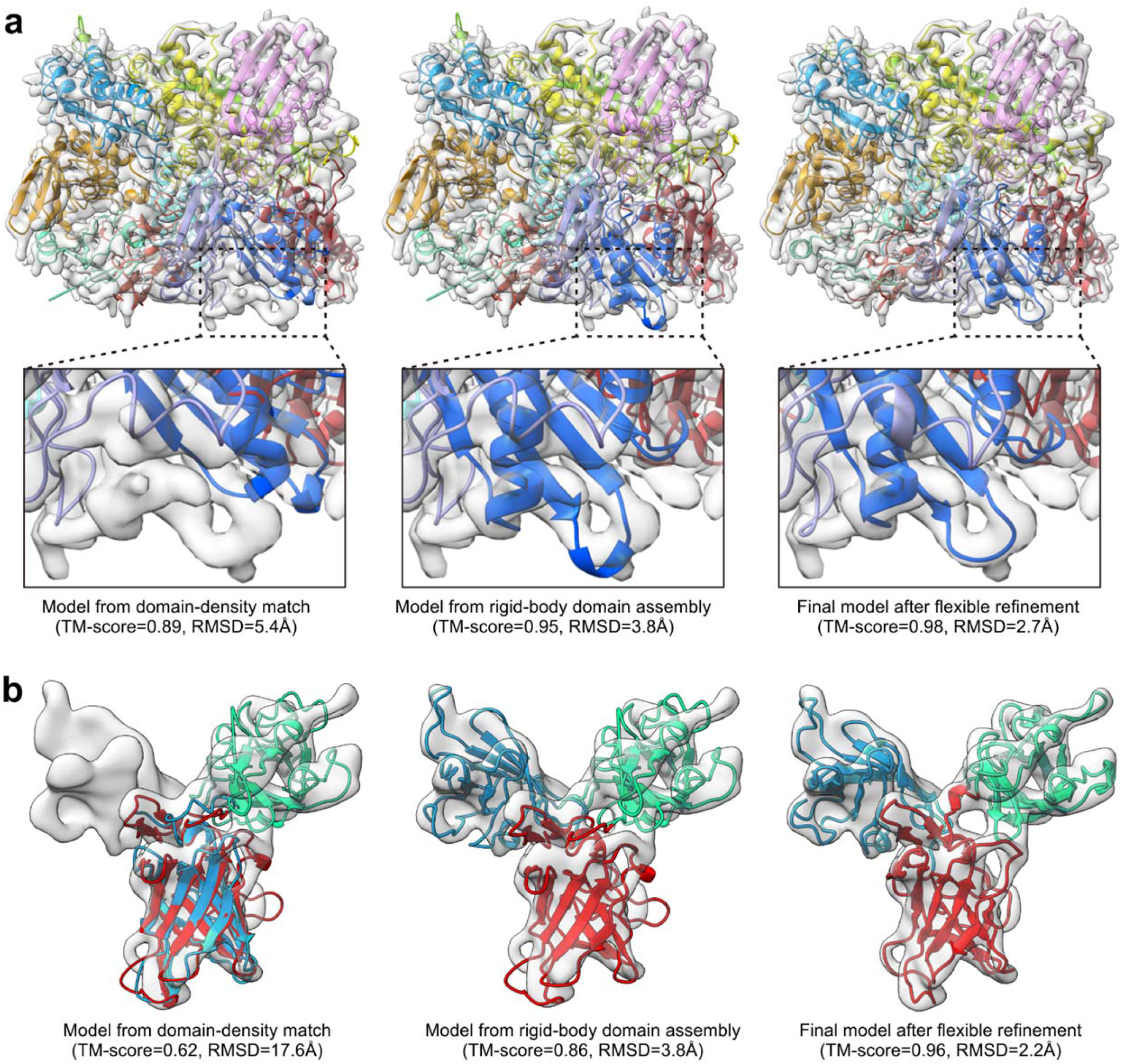
Representative examples showing the process of DEMO-EM. Density maps are shown in gray shadow where cartoons are DEMO-EM models with different colors indicating different domains. (**a**) 2e1qC, a protein with 10 domains (8 continuous domains and 2 discontinuous domains) using a simulated density map with a resolution of 5.3 Å. (**b**) 1q25A, a protein with 3 domains using a simulated density map with a resolution of 9.9Å.

**Figure 3b** shows another example from the *cation-independent mannose 6-phosphate receptor* (PDBID: 1q25A), a protein with 3 domains using a simulated density map with a low resolution of 9.9 Å. Two domains from N- and C-terminus were initially assigned into the same map space as they shared a similar fold with TM-score of 0.88; this resulted in a low TM-score=0.62 of full-length model. Guided by the global model-density correlation and particle movements implemented in the rigid-body assembly step, the incorrect domain fit was corrected with the full-length TM-score improved to 0.86. After the flexible assembly and refinement simulations, almost all wrong folding regions in the model were corrected which resulted in a global TM-score=0.96 and RMSD=2.2 Å. Again, the average TM-score/RMSD of the individual domains were significantly improved in this example from 0.76/3.6Å to 0.94/1.2Å, respectively.

In **Supplementary Figure 5**, we also present the full-length models created by MDFF and Rosetta starting with Situs for these two examples, which have a TM-score/RMSD equal to 0.17/34.5Å and 0.09/83.2Å for 2e1qC, and 0.36/22.2Å and 0.36/21.2Å for 1q25A, respectively; the low quality models are mainly due to the initial full-length models with incorrect domain orientations. The results of these case studies reinforce the advantage of the DEMO-EM pipeline for assembling multi-domain protein complex structures.

### Assemble multi-domain structures from experimental density maps

#### Dataset collection

To further examine the use of DEMO-EM on practical density maps, we collected a set of 51 non-redundant multi-domain proteins from EMDB that have experimental density map with resolution ranging from 2.8 to 10 Å (**Supplementary Fig. 6a**). The size of these proteins runs from 144 to 1,664 residues with the number of domains ranging from 2 to 8. To emulate the common real-life scenarios where the domain structures of target proteins are unknown, we predict the domain boundaries from sequence by a deep-learning contact-based program FUpred^14^, with the individual domain structures modelled by I-TASSER.

#### Overall benchmark results

As shown in **Figure 4a**, FUpred did an acceptable job in domain boundary prediction and correctly predicted the number of domains in 37 out of the 51 test proteins. In 70.6% of cases, the domain overlap rate is >80% compared to the DomainParser assignment on the target structure, resulting in an average normalized domain overlap (NDO) score of 0.83. The average TM-score of these domain models by I-TASSER was 0.72, where 94.0% of them had a correct fold with TM-score >0.5 (**Supplementary Fig. 6b**). After the DEMO-EM assembly, the full-length models have an average TM-score of 0.81 (**Fig. 4b**), where 92.2% of the cases have a correct global fold (TM-score>0.5). Meanwhile, the average TM-score of individual domains in the full-length models was increased from 0.72 to 0.77 with 85.9% domains improved (**Fig. 4c**), demonstrating again the ability of DEMO-EM on both levels of domain and full-length structure refinements.

**Figure 4.**
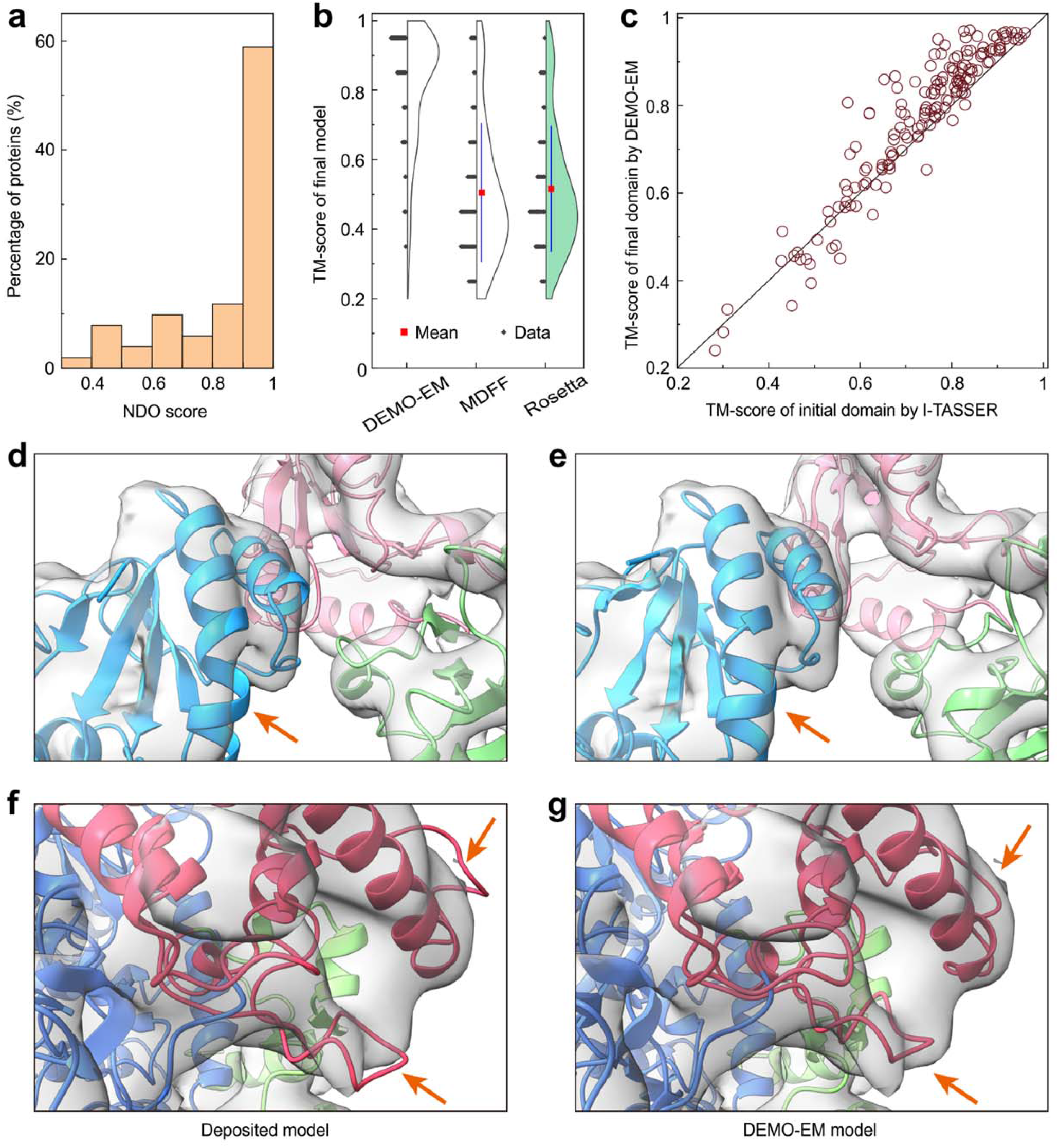
Summary of structures constructed by different approaches using experimental density map. (**a**) Distribution of NDO score of domain boundaries predicted by FUpred. (**b**) TM-score of full-length models constructed by DEMO-EM, MDFF, and Rosetta. (**c**) Head-to-head TM-score comparison of the initial individual models by I-TASSER and that in final full-length models by DEMO-EM. (**d, e**) The deposited model in PDB (PDBID: 6eny) (d) and reconstructed model (e) by DEMO-EM for *human PLC editing module*, where different color represents different domains. (**f, g**) The deposited model in PDB (PDBID: 5fj6) (f) and reproduced model (g) by DEMO-EM for *the P2 polymerase inside in vitro assembled bacteriophage phi6 polymerase complex*.

The average full-length TM-score (0.81) is slightly lower than that of the benchmark results on synthesized density maps (0.85), which is probably due to the fact that this dataset contains all targets while the former benchmark excluded targets with I-TASSER domains with incorrect folds. In fact, there are 10 out of 51 cases that have at least one I-TASSER domain model with TM-score <0.5, where the incorrect domain structures can result in incorrect domain-map match and misguide the subsequent domain assembly and refinement simulations. If we excluded the 10 cases with incorrect domain model, the overall TM-score increases to 0.86, which is largely consistent with the former benchmark data; this result also demonstrates the robustness of DEMO-EM whose performance does not depend on the source of density maps from syntheses or experiments.

As a control, we also listed in **Table 1** the modeling results by MDFF and Rosetta starting from the same set of I-TASSER domain models with the initial conformation assembled with Situs. The data shows again that DEMO-EM outperformed the control methods, with the average TM-score of the full-length models 76.1% and 102.5% higher than that of MDFF and Rosetta. The results are further confirmed by head-to-head TM-score comparison in **Supplementary Fig. 7**, where DEMO-EM has a higher TM-score for nearly all the targets.

#### Case studies showing DEMO-EM models are likely closer to native than deposited models

In Figures 4d-g, we present two representative examples with density maps taken from EMDB, in which the models generated by DEMO-EM are likely better than the structural models deposited in the databases. First, **Figure 4d** shows the deposited model of *human PLC editing module* in the PDB (PDBID: 6enyD with density map from EMD-3906 in EMDB), which was created by fitting of a homology model (from PDBID 3f8uC) with the cryo-EM density map at a resolution of 5.8 Å^27^ using FlexEM^8^ and Chimera^28^. Although the deposited model has a close similarity to the DEMO-EM model (**Fig. 4e**) with TM-score=0.96 and RMSD=1.4Å, many regions of the deposited model are exposed to the outside of the density map (e.g., the helix pointed by the arrow in **Fig. 4d**), which resulted in an overall density-model correlation coefficient (CC) of 0.82. Almost all these wrong regions were corrected in the model constructed by DEMO-EM with the CC value improved to 0.85 where the fitting of the entire DEMO-EM model is shown in **Supplementary Fig. 8a**.

**Figure 4f** shows the deposited model of another example from the *P2 polymerase inside in vitro assembled bacteriophage phi6 polymerase complex* (PDB 5fj6A with density map from EMD-3186), which contains 2 continuous domains mediated with a discontinuous domain. The deposited model was produced by the fitting of a homology structure (PDB 1hhsA) with the density map at a resolution of 7.9 Å^29^ using Chimera^28^ and Phenix^39^. As there are many significant noisy grid points in the density map, some regions of the deposited model (e.g. loops point by arrows in **Fig. 4f**) were incorrectly modelled because they were not wrapped in the density map with an overall CC of 0.83. The model created by DEMO-EM (**Fig. 4g**) on the same density map data fixed all these local errors and resulted in an improved CC score (=0.86). Again, while the overall structure of deposited and DEMO-EM models is largely consistent (TM-score=0.95 and RMSD=1.9 Å), the DEMO-EM model is likely closer to the target structure, based on the CC values and the better structure packing with the experimental density maps (see the overall fitting of the DEMO-EM model in **Supplementary Fig. 8b**).

Despite the favorable CC score values and better local structure packing, it is important to note that in the absence of other experimental information, it is still difficult to objectively assess if the DEMO-EM models are indeed closer to the native than that by other methods. For further examination, we made publicly available the DEMO-EM models for all the test proteins at https://zhanglab.ccmb.med.umich.edu/DEMO-EM/data_set/model_benchmark51.tar.gz.

### Application to structural modeling of the SARS-CoV-2 genome

Given the on-going COVID-19 pandemic, considerable effort has been made on structural determination of critical proteins in the SARS-CoV-2, a pathogen coronavirus having caused the pandemic^31^. In **Figure 5**, we present the full-length structural models constructed by DEMO-EM for all six SAR-CoV-2 proteins which have the cryo-EM data deposited in EMDB. Based on FUpred^14^ prediction, five proteins contain multiple domains and two of them include discontinuous domains (**Supplementary Table 4**). Compared to the deposited models, many of which contained missed residues due to the loss of density data, the DEMO-EM models have an average TM-score/RMSD of 0.97/1.4Å on the regions where the deposited models have structure. Compared to the original density maps, DEMO-EM models achieved a slightly higher average CC (0.82) than the deposited models (0.80).

**Figure 5.**
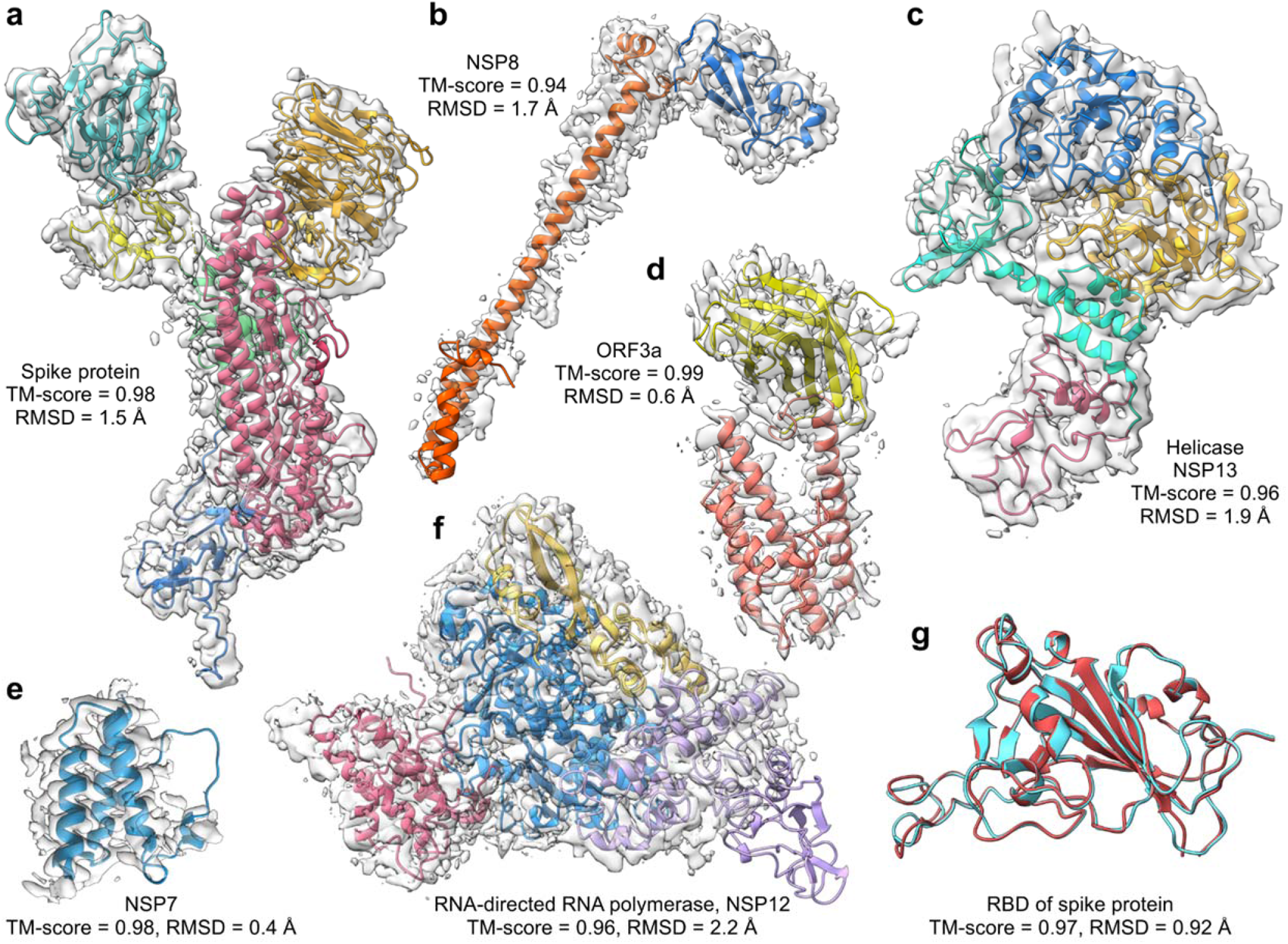
Overlay of structural models by DEMO-EM on the cryo-EM density maps for the six proteins in SARS-CoV-2 genome. (**a**) Spike protein (density map from EMD-21375). (**b**) NSP8 (EMD-11007). (**c**) Helicase/NSP13 (EMD-22160). (**d**) ORF3a (EMD-22136). (**e**) NSP7 (EMD-11007). (**f**) RNA-directed RNA polymerase/NSP12 (EMD-11007). (**g**) Comparison of the Spike RBD domain by DEMO-EM (cyan) with the X-ray structure (red, PDB 7bz5A).

The X-ray structure of the receptor binding domain (RBD) of the spike protein. which SARS-CoV-2 uses to bind the angiotensin-converting enzyme 2 (ACE2) to invade the host cells, was recently released (PDB 7bz5A)^32^. In **Fig 5g**, we show a comparison of the structural model built by the DEMO-EM with the released X-ray structure, where the DEMO-EM model has a TM-score of 0.97 and RMSD of 0.92 Å which are slightly better than that of the deposited model (TM-score=0.96 and RMSD =0.97 Å). The DEMO-EM models for all the six SARS-CoV-2 proteins are downloadable at https://zhanglab.ccmb.med.umich.edu/DEMO-EM/data_set/model_sarscov2.tar.gz.

## DISCUSSION

Due to the scarcity of multi-domain template structures in the PDB, automated determination of multi-domain protein structures from cryo-EM density becomes a significant challenge as most approaches in the community rely on fitting and refinement of homology models. To address this issue, we developed a new method, DEMO-EM, dedicated to structure assembly of multi-domain proteins from cryo-EM density maps. Without relying on global homologous templates, the method integrates cutting-edge single-domain modeling and deep residual network learning techniques with progressive rigid-body and flexible Monte Carlo simulations into a hierarchical pipeline that is ready for automated and large-scale multi-domain protein structure prediction.

DEMO-EM was carefully tested over a comprehensive benchmark set of 357 non-homologous proteins containing various numbers and types of domain structures using synthesized maps and 51 cases with experimental density data. The results showed that DEMO-EM yielded accurate full-length models for more than 91% cases with TM-score >0.8 using density maps ranging from 2-10 Å. Meanwhile, the accuracy of individual domains in the final full-length model was significantly improved with the average TM-score increased by 8.9%. DEMO-EM was controlled with two widely used cryo-EM based structure modeling methods (MDFF^8^ and Rosetta^13^). Starting from the same cryo-EM data and individually predicted domain models, the TM-score of the full-length models constructed by DEMO-EM is 32.5% and 39.5% higher than the control methods, corresponding to a p-value of 2.1E-98 and 1.1E-104 in Student’s t-test, respectively. As an application, DEMO-EM was used to model six proteins from the SARS-CoV-2 with cryo-EM data available and generated correct fold for all the targets, with some cases being likely closer to the native than the deposited models in the EMDB.

The superior performance of DEMO-EM stems partly from its ability for quick and reliably framework construction, which is enabled by the unique single-domain structure modeling from I-TASSER and the coarse-grained density-map space enumeration driven by the quasi-Newton searching process. Next, the domain-level rigid-body assembly simulation is capable of correcting domain positions and inter-domain orientations by combining density map restraints with inter-domain potentials, even when domain poses are occasionally incorrectly assigned in the initial frameworks. Finally, the atomic-level flexible structural assembly simulations couple density-map correlations with deep learning-based inter-domain distance profiles, which helps to simultaneously fine-tune local structural packing and inter-domain orientations and resulted in consistent improvement of both local and global structures.

Despite the promising domain assembly results, the applicability and accuracy of DEMO-EM could be further improved in several aspects. First, most of the density maps in our tests are segmented from the full density map by UCSF Chimera^28^. Although manual segmentation is often straightforward, the automatic map segmentation techniques (e.g. the method in *Phenix^30^)* can be introduced into DEMO-EM. Second, all individual domain models are directly produced by I-TASSER without using guidance from density data. An incorrect initial domain model may lead to a poor final model since it affects the algorithm to identify correct poses for initial framework constructions. Therefore, combining the restraints of density data with I-TASSER potentials for individual domain model generation will be helpful to improve the accuracy of final models. In addition, recent studies have shown the deep-learning based contact and distance maps can significantly improve the tertiary protein structure prediction accuracy^33,34^, an updated I-TASSER program combining the new neural-network based contact and distance restraints will further improve the quality of the DEMO-EM. Studies along these lines are under progress.

## ACKNOWLEDGMENTS

This work is supported in part by the National Institute of General Medical Sciences (GM136422 and S10OD026825 to Y.Z.), the National Institute of Allergy and Infectious Diseases (AI134678 to Y.Z.), the National Science Foundation (IIS1901191 and DBI2030790 to Y.Z.), the National Nature Science Foundation of China (61773346 to G.Z.) and the Key Project of Zhejiang Provincial Natural Science Foundation of China (LZ20F030002 to G.Z.). This work used the Extreme Science and Engineering Discovery Environment (XSEDE)^35^, which is supported by the National Science Foundation (ACI1548562).

## AUTHOR CONTRIBUTIONS

Y.Z. conceived the project. X.Z. developed the pipeline and performed the test. Y.L. developed the method for inter-domain distances prediction. W.Z. developed the method for domain boundaries prediction. C.Z. helped analyze the data. G.Z. helped supervise the research. X.Z. and Y.Z. wrote the manuscript, and all authors read and approved the final manuscript.

## COMPETING FINANCIAL INTERESTS

The authors declare no competing financial interests.

## METHODS

DEMO-EM is hierarchical approach to cryo-EM based multi-domain protein structure determination, which consists of four consecutive steps: (i) determining domain boundaries and modeling individual domains, (ii) matching domain models with density map for initial framework generation, (iii) rigid-body domain structure assembly for domain position and orientation optimization, and (iv) flexible structure assembly and refinement simulation of full-length structural models (**Fig. 1**).

### Domain parsing and individual domain structure folding

Starting from the query amino acid sequence, inter-residue contact maps are predicted by a deep-learning based neural network program, ResPRE^36^. The domain boundaries are then predicted using FUpred^14^ by maximizing the number of intra-domain contacts and minimizes the number of inter-domain contacts. Next, the structural model of each domain is generated using I-TASSER^15^ by assembling continuous fragments excised from threading templates identified from the PDB^37^. For discontinuous domains that contain 2 or more segments from separate regions of the query sequence, the domain models are obtained by sequentially connecting the sequences of all segments.

### Deep neural network-based inter-domain distance prediction

To help guide domain orientation assembly, an inter-domain distance map was predicted by a deep residual neural-network algorithm, DomainDist, whose architecture is outlined in **Supplementary Fig. 9**. DomainDist is an extension of the TripletRes^38^ that was originally developed to predict inter-residue contact maps based on a triplet of coevolutionary matrices but was extended here for predicting the probability of inter-residue distance within 36 bins in [2, 20] Å. The DomainDist program was trained on a non-redundant dataset of 26,151 proteins collected from the PDB, where the multiple sequence alignment (MSA) for each protein was constructed using HHblits^39^ searching against the Uniclust30 sequence database^40^. In addition to the 2D coevolutionary features employed in TripletRes, three 1D features, including Hidden Markov Model (HMM), one-hot representation of sequence and field parameters of Potts model, were adopted and tiled to two-dimension and concatenated with the 2D coevolutionary features. The neural network structure was designed following convolutional strategies, using ResNet basic blocks^41^. The neural network model was trained by Adam optimization algorithm to marginally minimize cross-entropy loss. Both intra-domain and inter-domain distance information was considered during the training, although only inter-domain distance information was considered by DEMO-EM.

### Quasi-Newton based domain and cryo-EM density-map matching

For each individual domain model from I-TASSER, we used Limited-memory Broyden–Fletcher–Goldfarb–Shanno (L-BFGS), a quasi-Newton optimization algorithm with 6D translation-rotation degrees of freedom, to identify the best location and orientation of the domain with the highest correlation with the density map (**Supplementary Fig. 10a**). Since L-BEGS is a local optimization method whose results depend on its initial solutions, we started L-BEGS simulation from multiple initial positions (translation vector) and orientations (rotation angle) by enumerating all combinations of Euler angles (∅, *θ*, and *ψ*) with a step size of 30° across the density-map space (**Supplementary Fig. 10b**). For a given domain pose, a density correlation score (DCS) calculated by

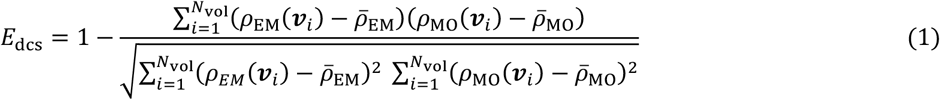

is used to guide the L-BEGS simulations. Here, *N*_vol_ is the number of voxels (grid points) in the density map and *ρ*_EM_(***ν**_i_*) is the experimental density of the *i*-th voxel ***ν**_i_*. The density probed from the decoy structure is calculated by 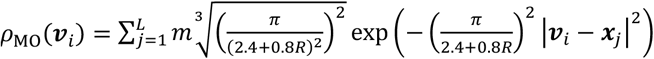, where ***x**_j_* is the position of the *j*-th atom in the decoy, *m* is its mass, and *R* is the resolution of the density map. To speed up the matching process, a density map with voxel size of 2Å interpolated from the original density map is employed. After the L-BEGS simulation, all poses for each domain with DCS <0.5 (or the top 10 poses when more than 10 poses with DCS<0.5) are pooled and combined with the top poses of other domains to form initial models of full-length models. The combination is made by permutating initial poses of all domains and allows for domain overlaps, where top 30 full-length models with the lowest DCS are selected for the next step of rigid-body domain match and assembly.

### Rigid-body domain assembly

Two rounds of rigid-body domain assembly simulations are performed to optimize domain positions and orientations. In the first round, domains are treated as particles where a quick replica-exchange Monte Carlo (REMC) simulation is made to quickly adjust the individual domain positions based on the global model-density correlations. In this step, the energy function contains only DCS (Eq. 1), where movements include rigid-body translation and rotation around each domain’s center of mass (**Supplementary Figs. 11a and b**). It is noted that here DCS is calculated for the full chain model which should lead to more optimal model result from the last step whose optimization was based on DCS from individual domains. The density map with a voxel size of 3Å interpolated from the original map is applied to reduce the computational cost. 30 replicas are sampled in parallel with the temperature ranging from 0.1 to 15, and a global swap movement between two neighboring replicas is performed for every 200 MC movements. The simulation is terminated when the number of swaps reaches 20 ∗ *N*_dom_. where *N*_dom_ is the number of domains. The top 30 models are selected according to the DCS for the next round.

The second round of rigid-body REMC simulation is to fine-tune the domain poses with a more detailed energy force field as defined in Eqs. (2-5), where a more elaborate density map with a voxel size of 2 Å is interpolated from the original density map for the assembly. Besides the translation and rotation movements used in the first round, three new movements are added (**Supplementary Figs. 11c-e**), including self-rotation around the N-to-C axis of each domain, the translation along the neighboring domains in sequence, and the pose exchange between two domains with similar structures (i.e., TM-score ≥ 0.75) according to TM-align^42^, which is designed to reduce the case where domains with similar topology are swapped in the initial positions. The similar parameter setting with the first round is employed for REMC simulation, but the top 40 models are selected according to DCS for the next step.

### Energy function for rigid-body simulation

Conformations in the rigid-body assembly are assessed by an energy function with four terms:

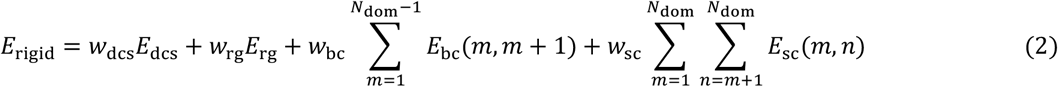

where the first term *density correlation score* is the same as defined in Eq. (1) but here it is used for full-length model.

The second term is *radius of gyration restraint*, defined as

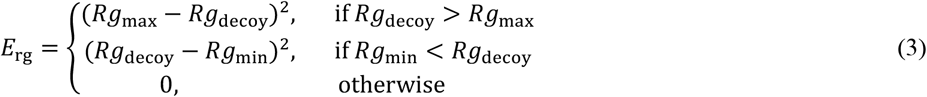

where *R*_*g*_decoy__ is the radius of gyration of the decoy structure, *R*_*g*_max__ and *R*_*g*_min__ are the maximum and minimum estimated radius of gyration, respectively, i.e., 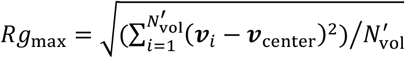, which is calculated from the 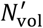 voxels with density ≥0.05 after normalizing density values to the range of 0 and 1, where 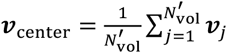 is the center point of these voxels. *R*_*g*_min__ = 2.849*L*^0.319^ (*L* is the query sequence length) is the statistical radius of gyration based on the known multi-domain protein models in the PDB, which has a Pearson correlation coefficient of 0.995 with real values (**Supplementary Fig. 12**).

The third term *domain boundary connectivity* is designed to constrain the connectivity of two neighboring domains along sequence (*m < n*), which is calculated by

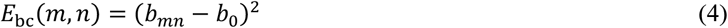

where *b_mn_* is the *C_α_* atom distance between the C-terminal residue of the *m*-th domain and the N-terminal of the *n*-th domain. For the case including discontinuous domains, *b_mn_* = (*d*_1_ + *d*_2_)/2 is the average of two linker distances connecting the continuous domain with the discontinuous segments (see **Supplementary Fig. 13**). *b*_0_ = 3.8 Å is the standard distance between neighboring Cα atoms.

The last term *steric clashes* penalizes domain pairs occupying the same space, which is defined as

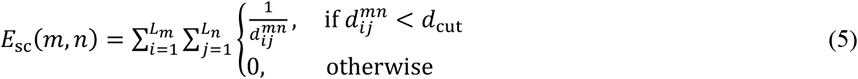

where *L_m_* and *L_n_* represent the sequence length of the *m*-th and *n*-th domain, respectively. 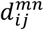 is the distance between the *i*-th *C_α_* atom of the *m*-th domain and the *j*-th *C_α_* atom of the *n*-th domain in the decoy structure. *d*_cut_ = 3.75 Å is the cutoff of distance to define a clash.

The weighting factors in *E*_rigid_ is optimized based on a training set of 425 proteins that has sequence identity <30% to the test proteins, by maximizing the correlation between total energy and RMSD of the decoy models to the native using the differential evolution algorithm^43,44^. This resulted in *w*_dcs_ = 300, *w*_rg_ = 1.13, *w*_bc_ = 0.55, and *w*_sc_ = 0.91.

### Atom-level flexible domain assembly and refinement

The process of flexible domain assembly and refinement contains two stages of simulations with progressive voxel resolutions and sampling focuses. In the first stage, six different movements are implemented (**Supplementary Fig. 14**): (i) LMProt^45^ perturbation; (ii) segment rotation around the axis connecting two terminus; (iii) conformational shift of segments along the sequence; (iv) rigid-body segment translation; (v) rigid-body tail rotation; (vi) rigid-body domain-level translation and rotation. To enhance the efficiency, a nine-residue sliding window is used to determine which region needs more aggressive conformation sampling, where a local score (*LC_i_*) for the sliding window of the center residue (*i*) is computed as the average correlation coefficient between the nine-residue fragment and the entire density map. The probability for the *i*-th residue to be selected for movement is set as

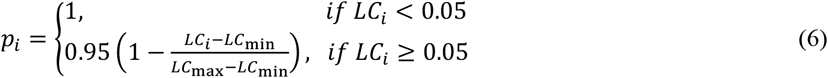

where *LC*_max_ and *LC*_min_(= 0.05) represent the maximum and minimum local scores, respectively. As illustrated in **Supplementary Fig. 15**, the setting in Eq. (6) helps ensure that the residues poorly correlated with the density map can receive more sampling than others. An atomic-level force field (see Eqs. 7-13 below) is designed to guide the REMC simulation at this stage, where a density map with a voxel size of 3Å interpolated from the original density map is applied to reduce the computation cost and the DCS is calculated based on backbone atoms. Similarly, 40 replicas with the temperature ranging from 0.01 to 15 are sampled in parallel. The global swap movement between two neighboring replicas is performed for every 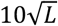 movements, where the simulation stops when the number of swaps reaches 200. All accepted decoys in the simulation are clustered by SPICKER^17^, and the centroid model in the first cluster is selected as a reference model for the second stage.

In the second stage, a finer density map with a voxel size of 2 Å is implemented with the DCS computed on all atoms. In addition, all residues have the equal probability to be selected for the movement and sampling. The REMC simulation is guided by the same force field defined in Eqs. (7-13) but the reference model in Eq. (10) is replaced by the centroid structure of the first cluster determined by SPICKER in the first stage. The simulation is terminated when the number of swaps reaches to 100. The lowest-energy decoy is selected to construct the final model, with the side-chain atoms repacked by FASPR^18^ followed by the FG-MD^19^ refinement.

### DEMO-EM force field for flexible assembly simulation

The flexible domain assemble simulations are implemented at a semi-atomic level, with each residue represented by N, *C_α_*, C, O, C_β_, H, and side-chain center of mass (SC). Among the seven modeling units, only the three backbone atoms (N, *C_α_*, and C) have coordinates directly determined in conformation sampling, while the other four are determined based on their relative positions to the three backbone atoms with parameters listed in **Supplementary Table 5**. The simulations are guided a composite force field consisting of seven energy terms:

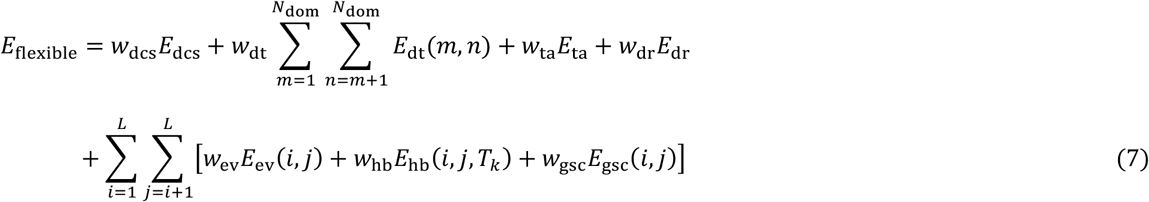

The first term counts for the *density correlation*, which is in the same form as Eq. (1) but calculated for full-length model. The second term is the *inter-domain C_β_ distance map* as predicted by DomainDist:

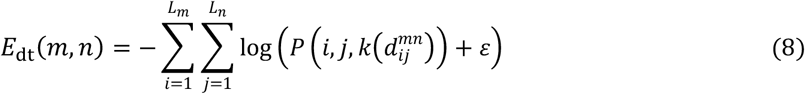

where 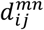 is the distance between the *i*-th *C*_β_ (*C_α_* for Glycine) atom in the *m*-th domain and *j*-th *C_β_* atom in the *n*-th domain, 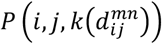 is the predicted probability of the distance 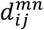 located in the *k*-th distance bin, and *ε* = 1*E* – 4 is the pseudo count to offset low-probability bins. In the calculation, we only consider atom pairs with probability peak located in [2Å, 20Å], and these atom pairs with predicted probabilities >0.5 in the last bin [>20 Å], which represents a low prediction confidence in [2Å, 20Å], are excluded.

The third term counts for *torsion angle variations* by

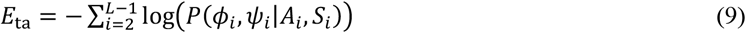

where *φ_i_* and *ψ_i_* represent the torsion angle pair of the *i*-th residue; *A_i_* and *S_i_* are the residue type and secondary structure type of the *i*-th residue, respectively; *P*(*φ_i_,ψ_i_*|*A_i_,S_i_*) is the conditional probability calculated based on the Ramachandran map of 6,023 high-resolution protein structures from the PDB^46^.

The fourth term *domain structure restraint* is to prevent topologies of individual domains deviating too far away from the initial structures generated by I-TASSER:

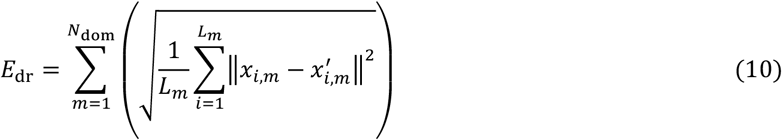

where *L_m_* is the sequence length of the *m*-th domain; *x_i,m_* represents the *i*-th *C_α_* atom in the *m*-th domain of the decoy after superposing the domain to the reference model by I-TASSER, and 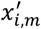 is the corresponding atom in the reference model.

The fifth term *excluded volume interaction* is defined as

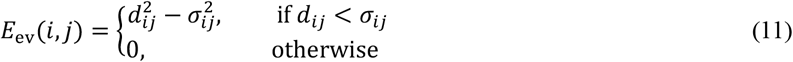

where *d_ij_* is the distance between the *i*-th and *j*-th atoms from different residues and *σ_ij_* is the sum of the van der Waals radius of the atom pairs taken from CHARMM^47^ (**Supplementary Table 6**).

The sixth term is the *hydrogen bonding* extended from QUARK^46^. As shown in **Supplementary Fig. 16**, only backbone H-bonds between residues (*i* and *j*) are considered, where four geometric features, including the distance between O_*i*_ and H_*j*_ (*D*(O_*i*_, H_*j*_)); the inner angel between C_*i*_, O_*i*_, and H_*j*_ (*A*(C_*i*_,O_*i*_,H_*j*_)); the inner angle between O_*i*_, H_*j*_, and N_*j*_ (*A*(O_*i*_,H_*j*_,N_*j*_)); and the torsion angle between C_*i*_, O_*i*_, H_*j*_, and N_*j*_ (*T*(C_*i*_,O_*i*_,H_*j*_,N_*j*_)), are selected to evaluate the bonding. We consider four types of Hydrogen bonds (*T*_1_: helix, *j = i* + 4; *T*_2_: helix, *j = i* + 3; *T*_3_: parallel β-sheets; and *T*_4_: antiparallel β-sheets). The energy term of a single backbone hydrogen bond is thus calculated by

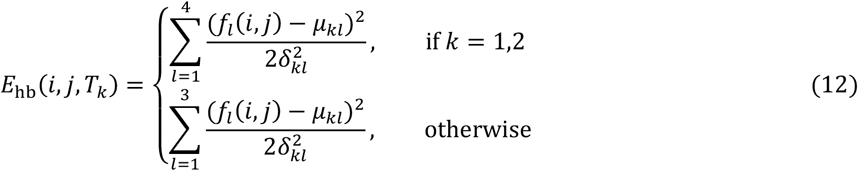

where *T_k_* represents the *k*th type of hydrogen bond, *f_l_*(*i,y*) denotes the *l*th feature of the decoy structure; *μ_kl_* and *δ_kl_* are the mean and standard deviation of the *l*th feature in the *k*th type hydrogen bond, which were pre-calculated from the high-resolution PDB structures and listed in **Supplementary Table 7**.

The last term *generic sidechain-atom contact potential* is used to evaluate the contacts between SC in one residue (*i*) and N, *C_α_*, C, O, C_β_ and SC atoms in another residue (*j*):

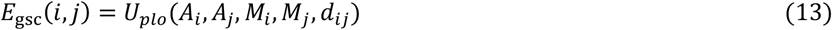

where *A_i_* (or *A_j_*) is amino acid type of residue *i* (or *j*), *M_i_* (or *M_j_*) represents the atom type of the *i* (or *j*)-th residue, *d_ij_* is the distance between the SC of the *i*-th residue and *M_j_* atom of the *j*-th residue, and *U_plo_*(*A_i_,A_j_,M_i_,M_j_,d_ij_*) is the corresponding polarity potential pre-calculated from 6,500 non-redundant high-resolution PDB structures (see https://zhanglab.ccmb.med.umich.edu/DEMO-EM/potential.html).

Similarly, the weighting parameters in Eq. (7) are determined by maximizing the correlation between the total energy and RMSD of the structure decoys of the 425 training proteins. This results in *w*_dcs_ = 320, *w*_ta_ = 0.3, *w*_dr_ = 1.5, *w*_dt_ = 0.15, *w*_ev_ = 0.1, *w*_hb_ = 0.05, and *w*_gsc_ = 0.1.

## Notes

### Competing Interest Statement

The authors have declared no competing interest.

