## Supplementary File for "Progressive and accurate assembly of multi-domain protein structures from cryo-EM density maps"

### Supplementary Information

#### Supplementary Figures

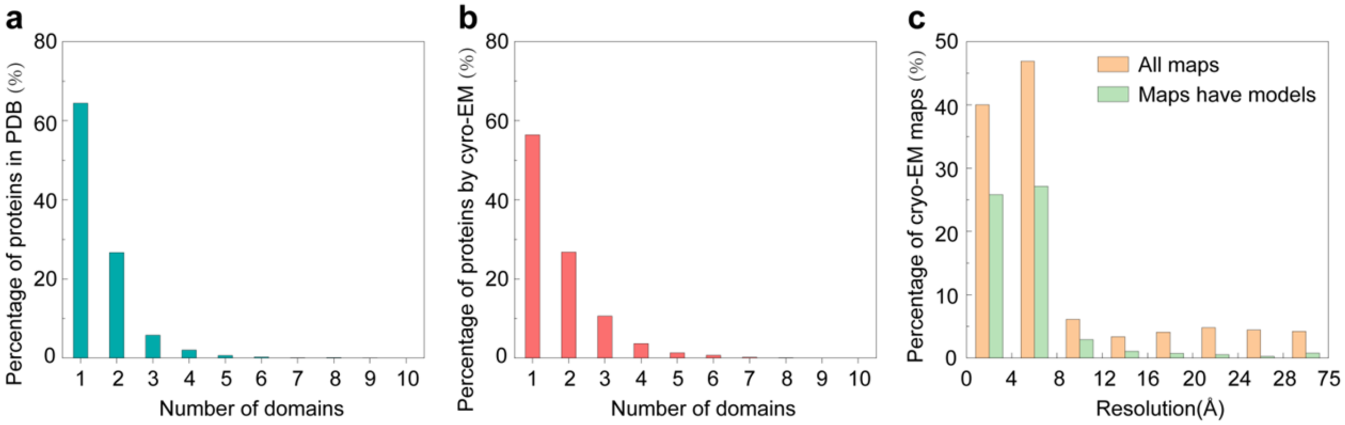


**Supplementary Figure 1**

The proportion of multi-domain proteins in protein chains in the PDB and the proportion of released cryo-EM maps with associated atomic models in EMDB.

(**a**) Distribution of multi-domain proteins in protein chains in the entire PDB, which indicates that only 35.6% proteins have more than 2 domains. (**b**) Distribution of multi-domain proteins in protein chains determined by cryo-EM, which shows that 43.6% cases are multi-domain proteins, where domain boundaries determined by CATH database[1] and DomainParser[2]. Here, we just show proteins with number of domains less than 10. (**c**) Distribution of resolutions for all released cryo-EM density maps and maps have corresponding atomic structures in EMDB. The data show that only 49.5% maps have associated atomic structures.

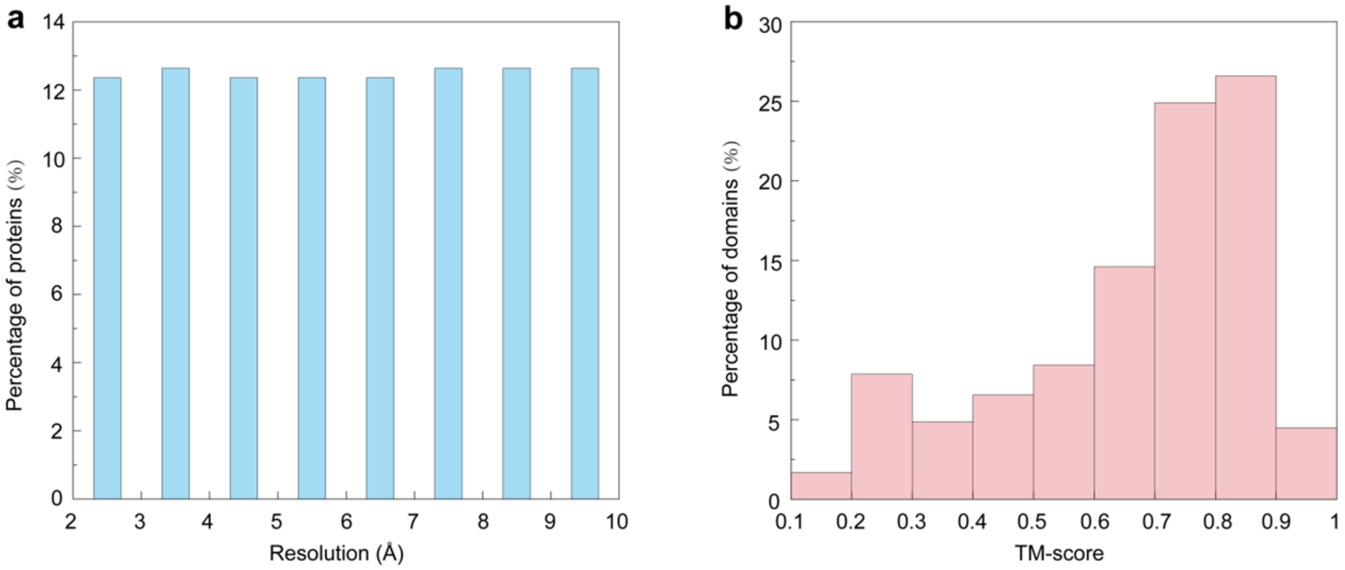


**Supplementary Figure 2**

Initial information of the 357 test multi-domain proteins

(**a**) Distribution of resolutions for simulated density maps. (**b**) Distribution of TM-scores for all domain models predicted by I-TASSER

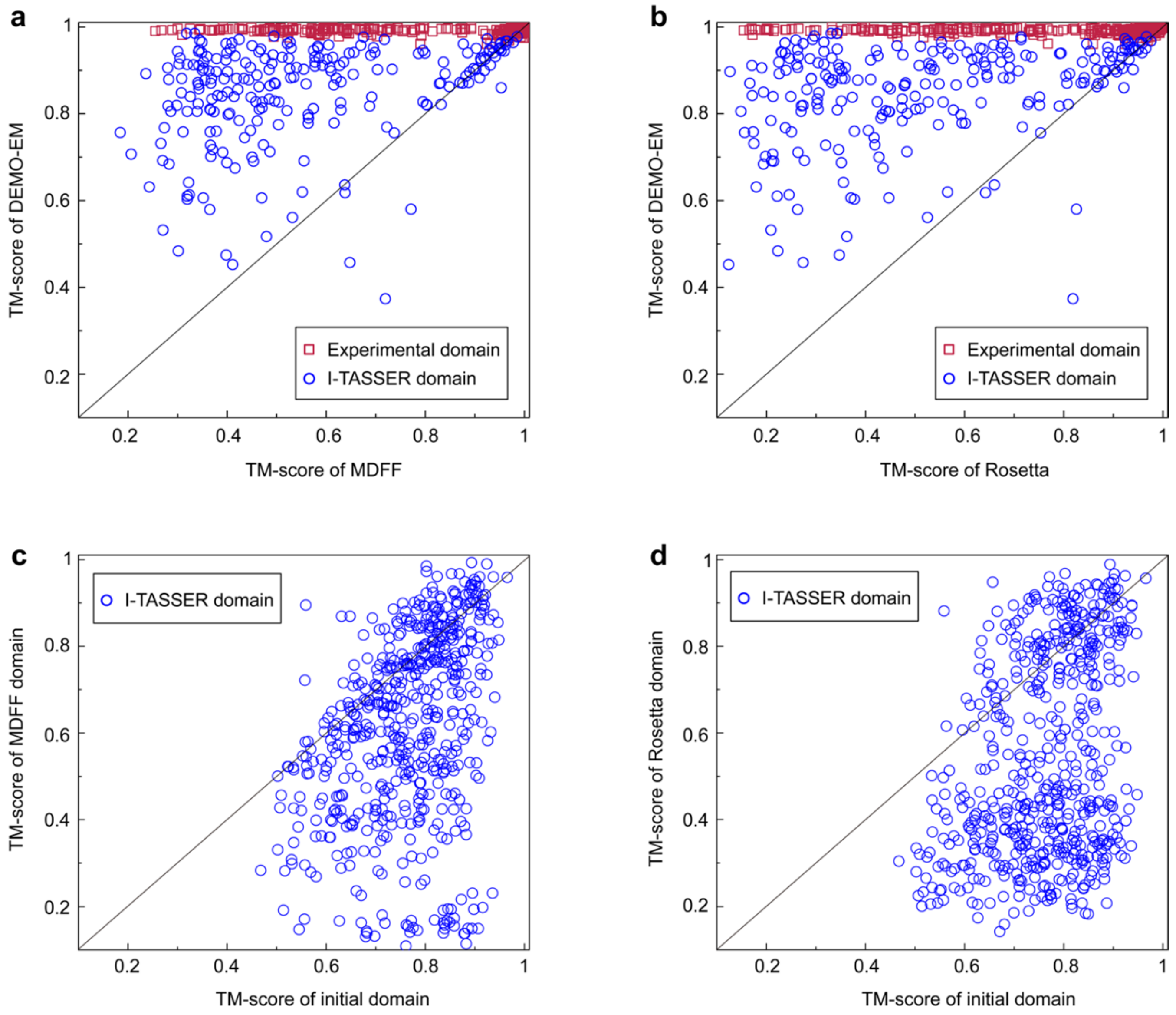


**Supplementary Figure 3**

Summary of models generated by MDFF[3] and Rosetta[4] using Situs[5] created initial full-length models and synthesized density maps for the 357 proteins.

(**a**) Head-to-head comparison between TM-scores of final models built by DEMO-EM and MDFF. (**b**) Head-to-head comparison between TM-scores of final models built by DEMO-EM and Rosetta. **(c)** Head-to-head comparison between TM-scores of initial individual domain models and the corresponding domain models in final full-length models by MDFF. **(d)** Head-to-head comparison between TM-scores of initial domain individual models and the corresponding domain models in final full-length models by Rosetta.

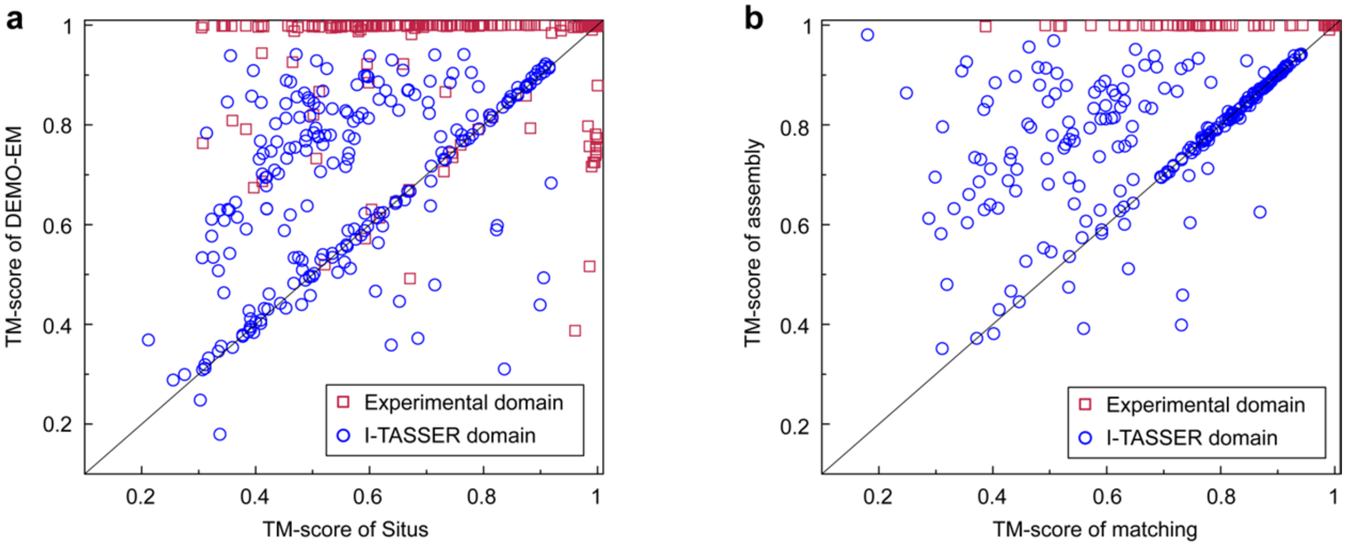


**Supplementary Figure 4**

Summary of models generated by domain matching and rigid-body assembly of DEMO-EM for the 357 test proteins.

(**a**) Head-to-head comparison between TM-scores of initial full-length models generated by independently fitting each domain into density maps using DEMO-EM versus that created by Situs using the same domain models. (**b**) Head-to-head comparison between TM-scores of initial full-length models from domain-map fitting versus that of rigid-body assembled models by DEMO-EM.

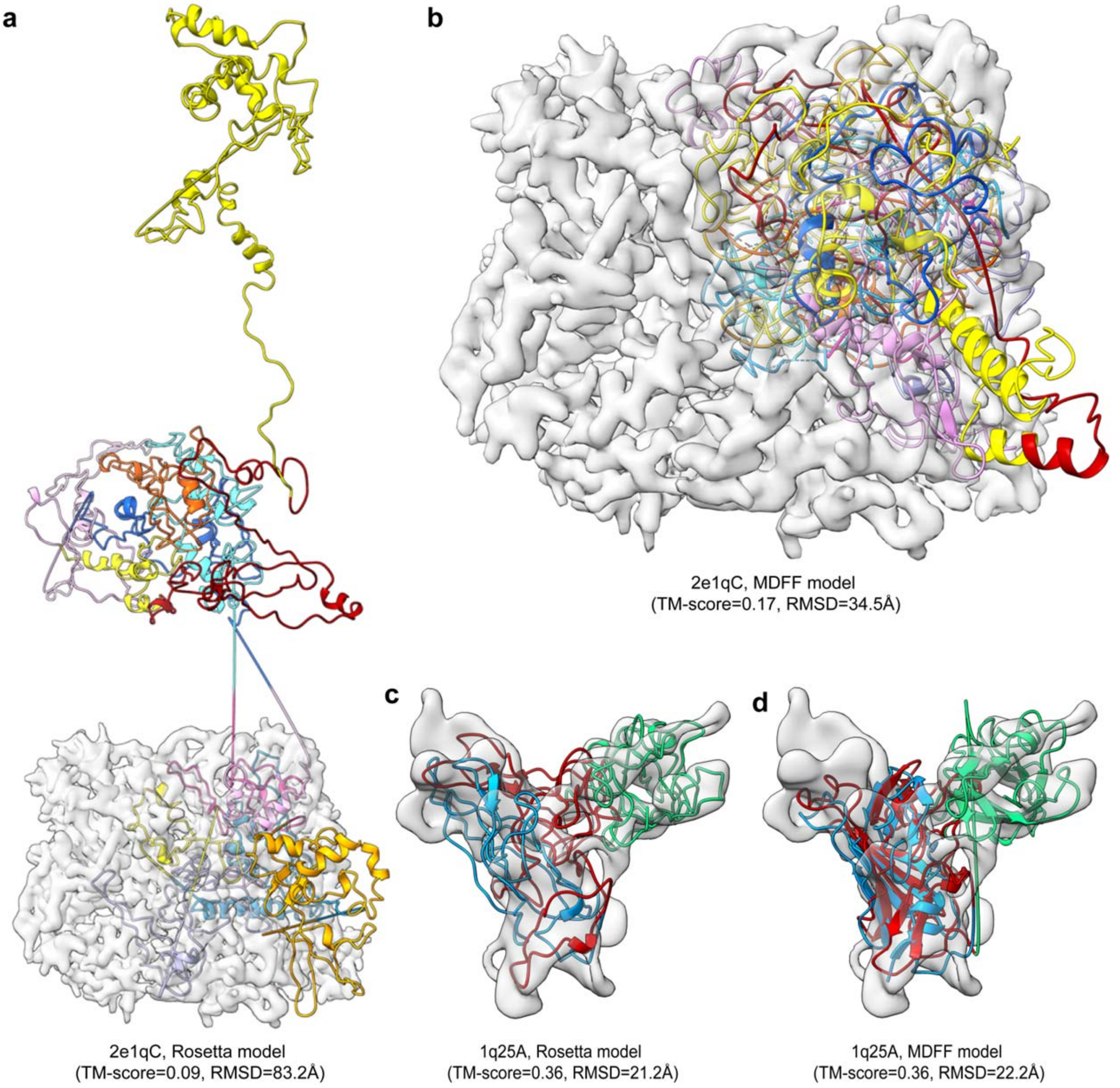


**Supplementary Figure 5**

Examples to show models refined by MDFF and Rosetta starting from full-length models generated by Situs.

(**a**), (**b**) Models of 2e1qC refined by Rosetta and MDFF, respectively. (**c**), (**d**) Models of 1q25A refined by Rosetta and MDFF, respectively.

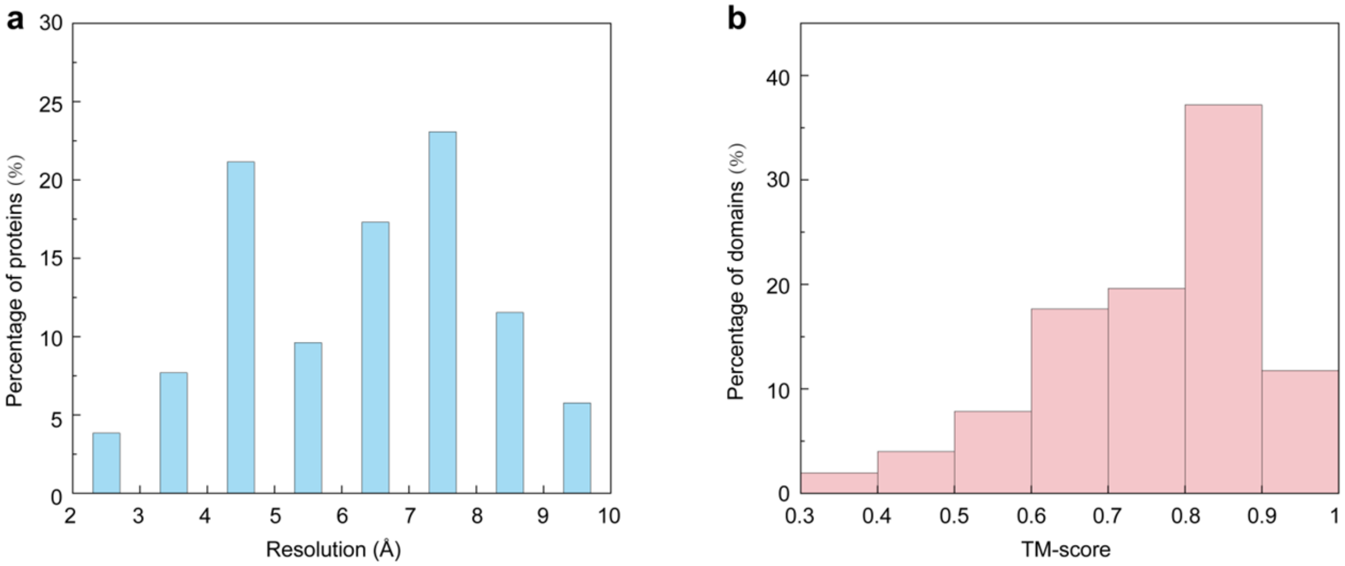


**Supplementary Figure 6**

Initial information for the 51 cases with experimental density maps.

**(a)** Distribution of resolutions for density maps. **(b)** Distribution of TM-scores for domain models generated by I-TASSER.

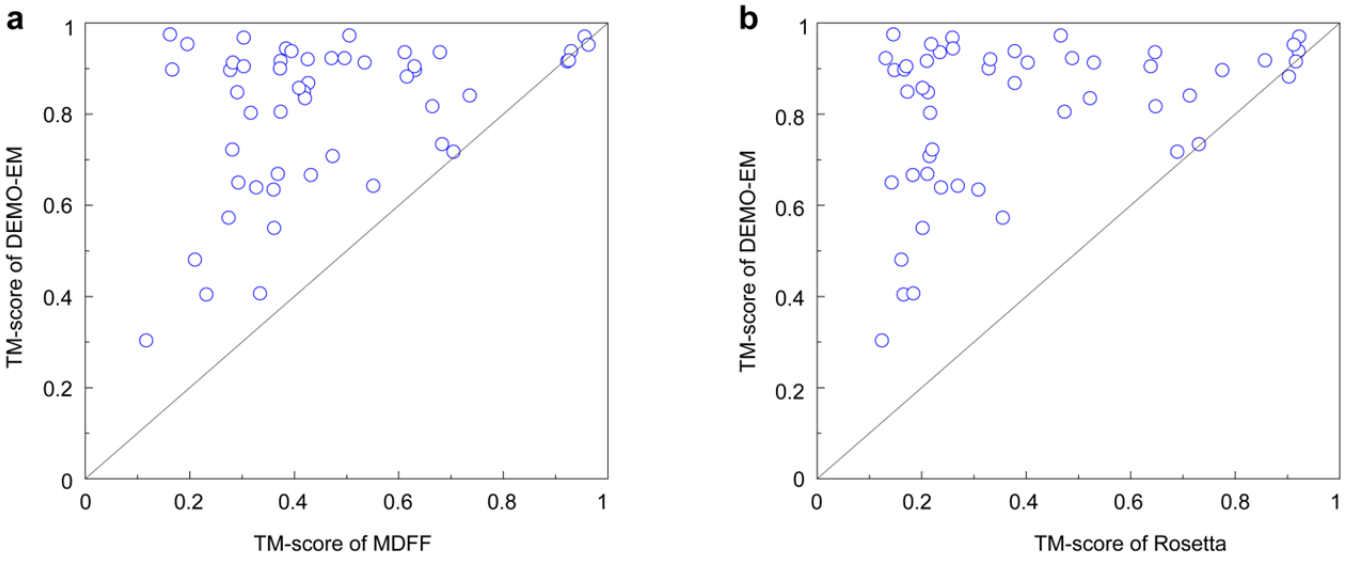


**Supplementary Figure 7**

Summary of models generated by different methods the 51 cases with real density maps.

**(a)** Head-to-head comparison between TM-scores of final model generated by DEMO-EM and that by MDFF using I-TASSER domain models. **(b)** Head-to-head comparison between TM-scores of final model and that by Rosetta using I-TASSER domain models.

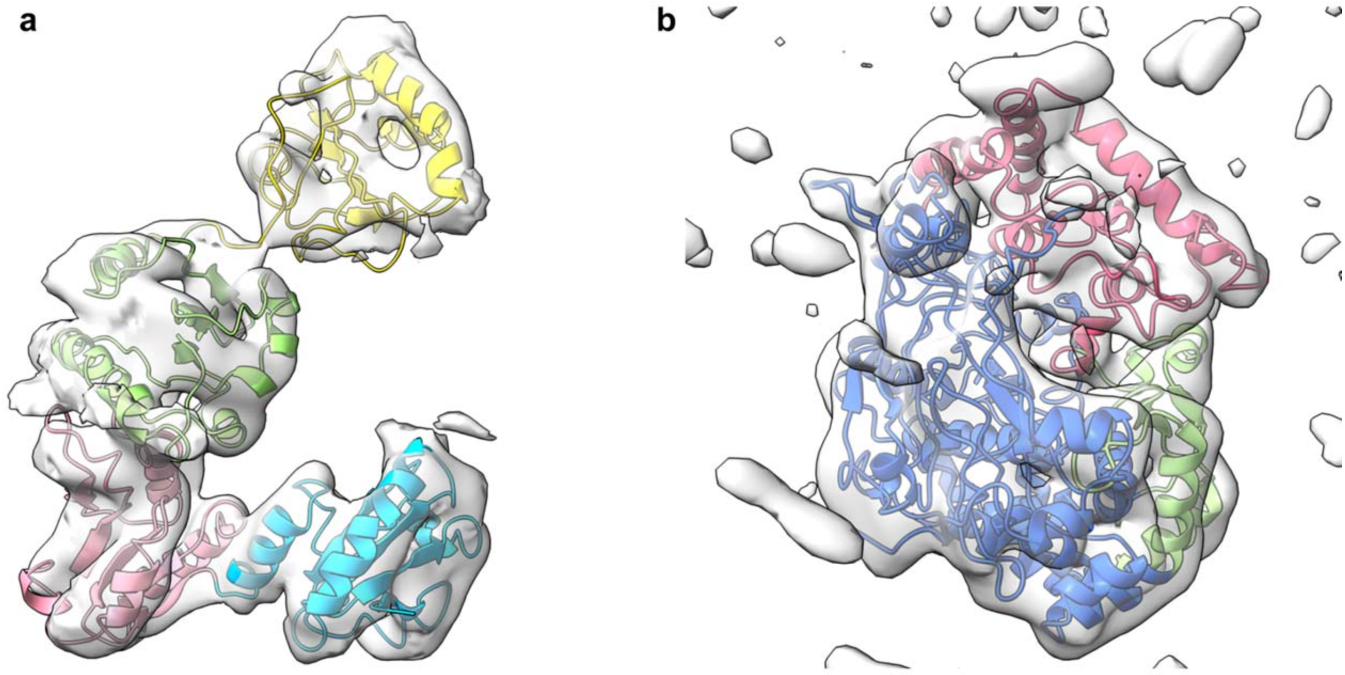


EMD-3906 (PDB code 6enyD), 5.8Å
TM-score = 0.96, RMSD = 1.4Å

EMD-3186 (PDB code 5fj6A), 7.9Å
TM-score = 0.95, RMSD = 1.9 Å

**Supplementary Figure 8**

Representative examples to show DEMO-EM constructed models using experimental density maps, where different domains represented by different colors.

(**a**) 6enyD (EMD entry 3906), a protein with 4 continuous domains. (**b**) 5fj6A, a protein with two continuous domains and one discontinuous domain (green).

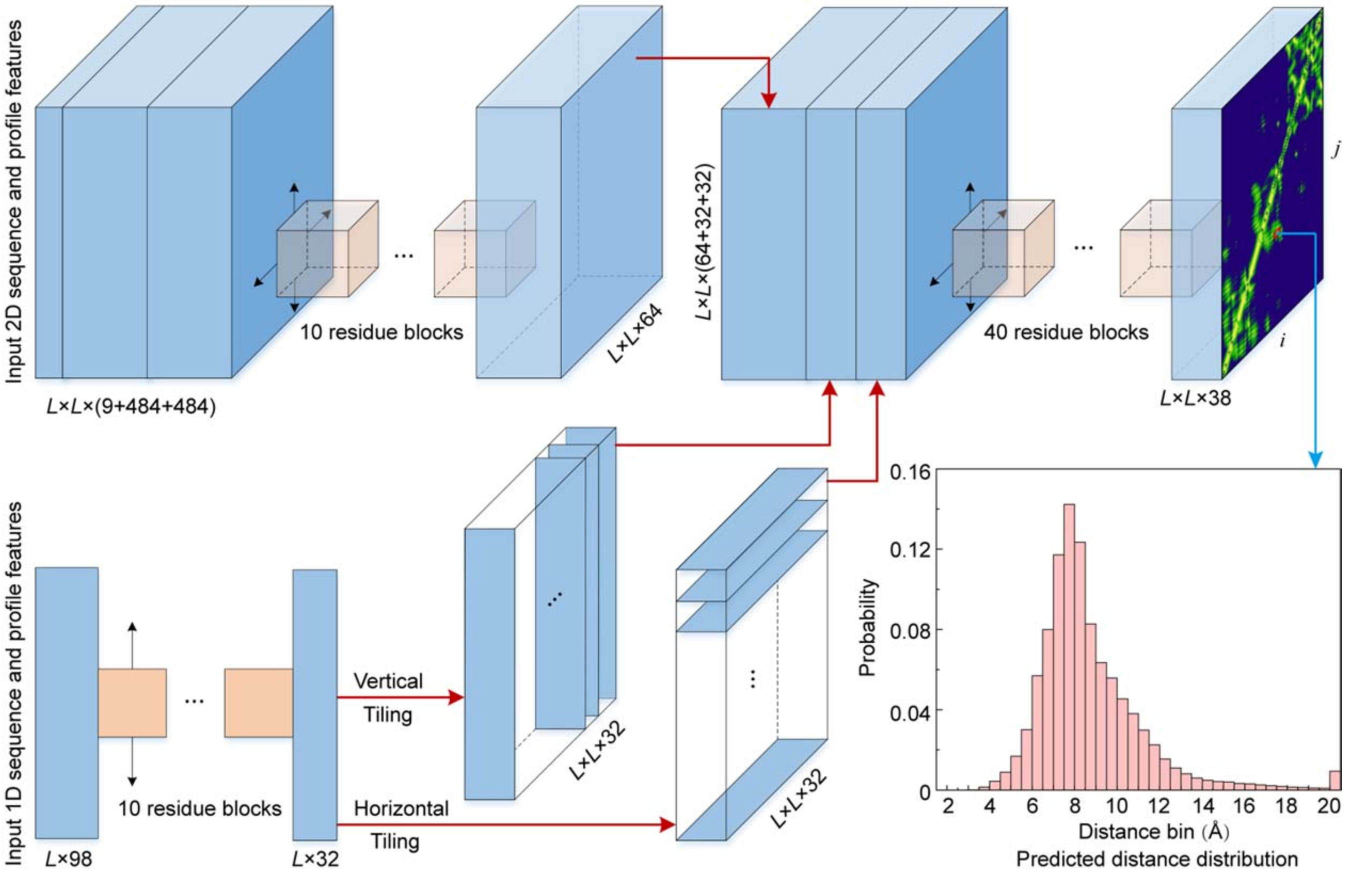


**Supplementary Figure 9**

Outline of DomainDist for predicting histograms of inter-domain and intra-domain residue-pair distances using deep-learning based neural networks.

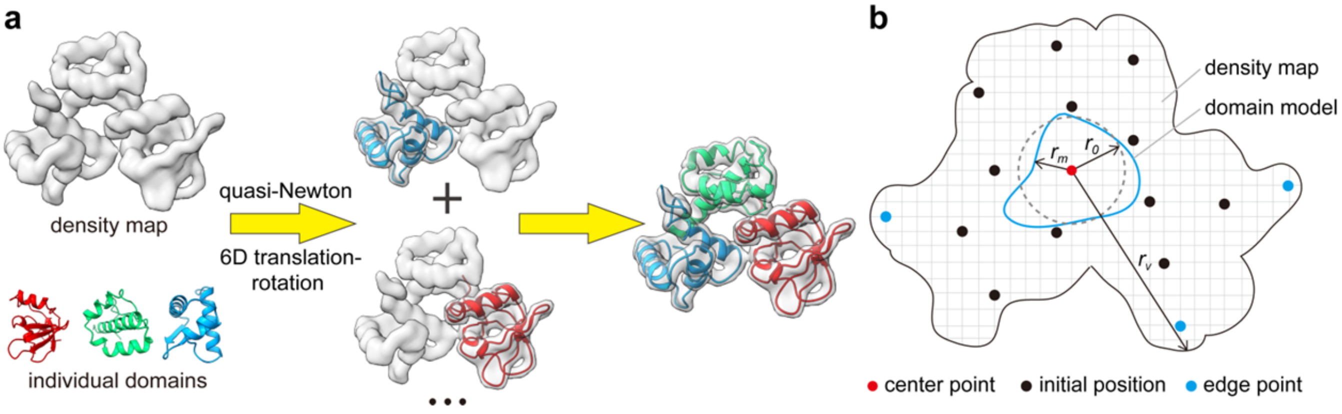


**Supplementary Figure 10**

Matching domain models into cryo-EM density maps

(**a**) The pipeline of domain structure matching into cryo-EM density maps in DEMO-EM. (**b**) Enumeration of initial position determination for a domain model. To eliminate redundant position, we set the minimum distance between two neighboring positions, $r = \max(0.5r_m, r_0)$, where $r_m$ is the radius of gyration of the model, and $r_0 = 5Å$ is the minimum distance between two initial positions. To remove the edge positions, the maximum distance between each initial position and the center point of the density map is set as $R_{\text{eg}} = \min(\max(1.1(r_v - r_m),\ r_0), r_v)$, where $r_v = \sqrt{(\sum_{i=1}^{N'_{\text{vol}}}(\boldsymbol{v}_i - \boldsymbol{v}_{\text{center}})^2)/N'_{\text{vol}}}$ is the radius of gyration of the density map calculated by the $N'_{\text{vol}}$ voxels with density ≥0.05 after normalizing density values to the range of 0 and 1, and $\boldsymbol{v}_{\text{center}} = \frac{1}{N'_{\text{vol}}}\sum_{j=1}^{N'_{\text{vol}}}\boldsymbol{v}_j$ is the center point of these voxels.

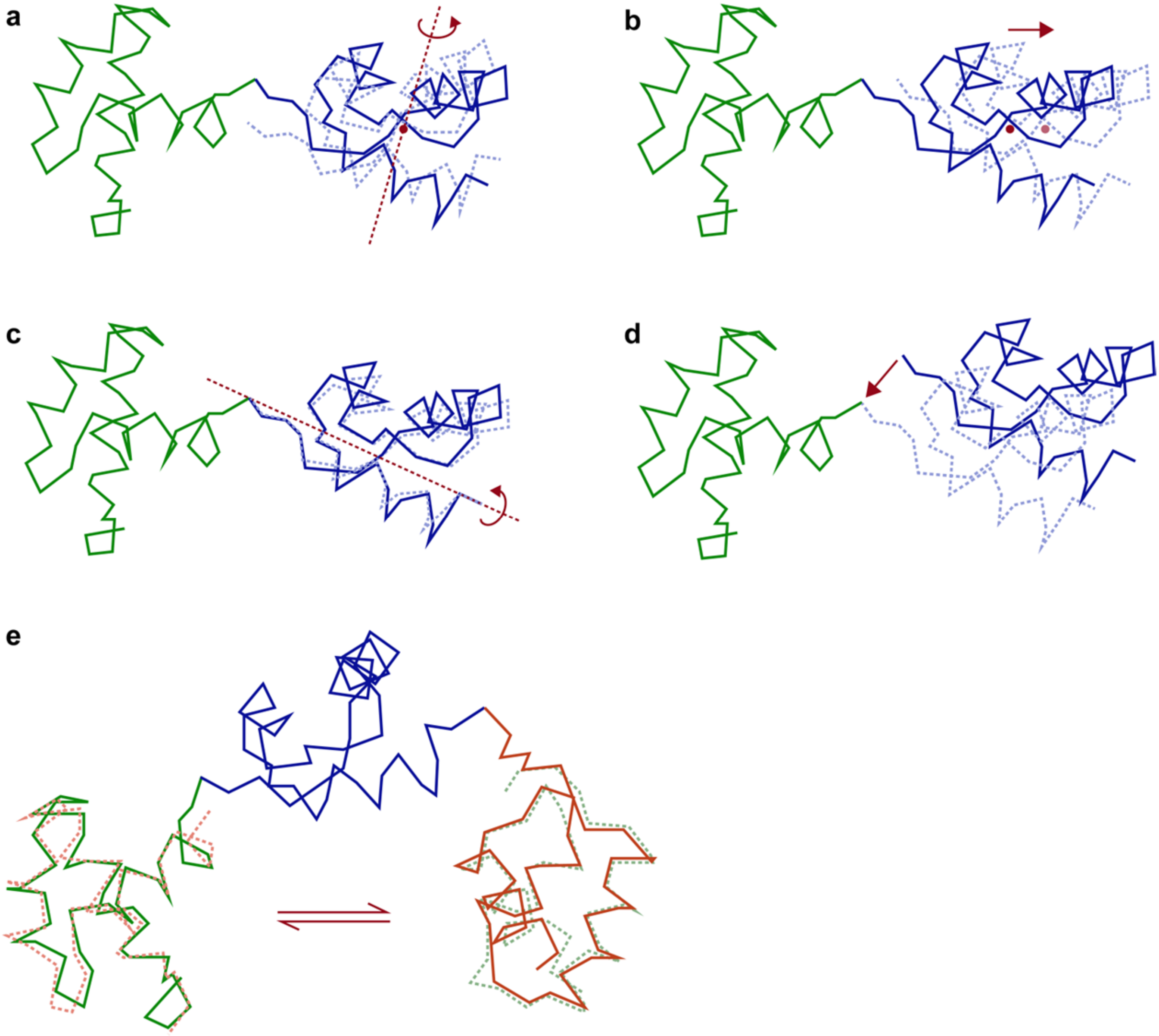


**Supplementary Figure 11**

Movements for the rigid-body domain assembly.

(**a**) Random rigid-body rotation around the domain's center of mass. (**b**) Random rigid-body translation of the domain's center of mass. (**c**) Random rigid-body rotation around the axis connecting the domain's N- and C-terminal $C_\alpha$ atoms. (**d**) Rigid-body translation along the axis connecting two domains which are neighboring in sequence. (**e**) Pose exchange between two domains with similar structures.

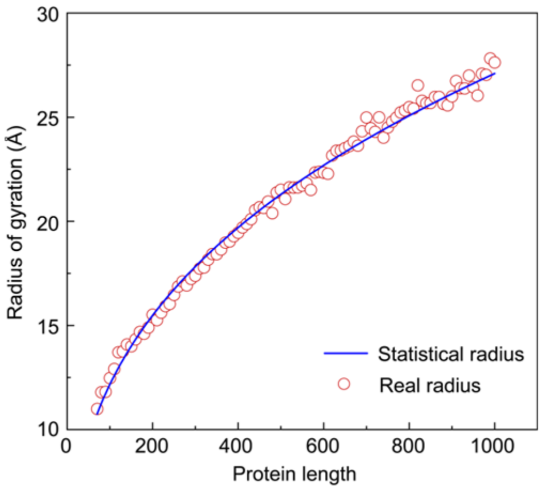


**Supplementary Figure 12**

Comparison the statistical minimum radius of gyration with the real values.

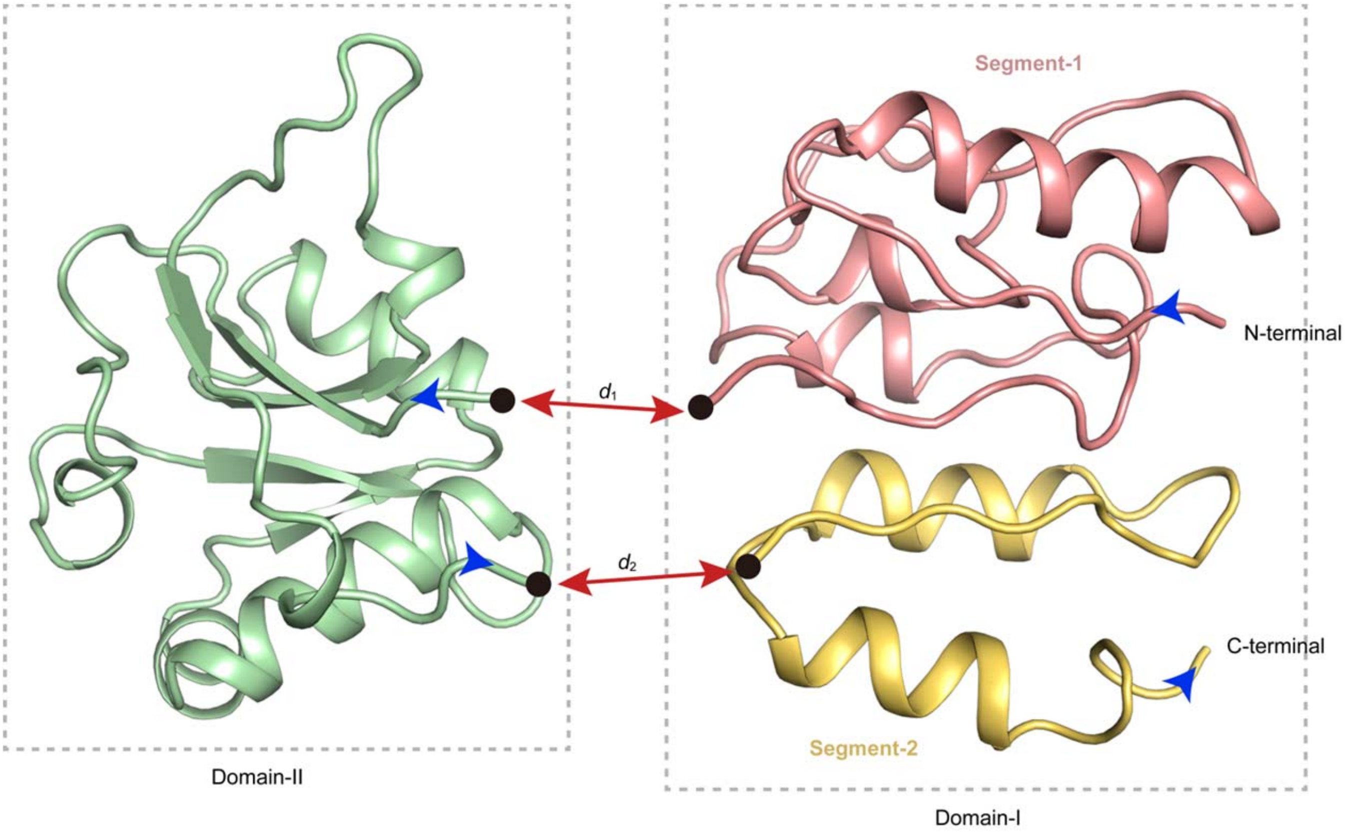


**Supplementary Figure 13**

Illustration of domain boundary distance potential for a two-domain protein with discontinuous domains. The discontinuous domain (Domain-I) is split into two segments due to the insertion of the continuous domain (Domain-II). $d_1$ and $d_2$ are $C_\alpha$-distances which are constrained to 3.8 Å by boundary distance energy (see Eq. (4) of method).

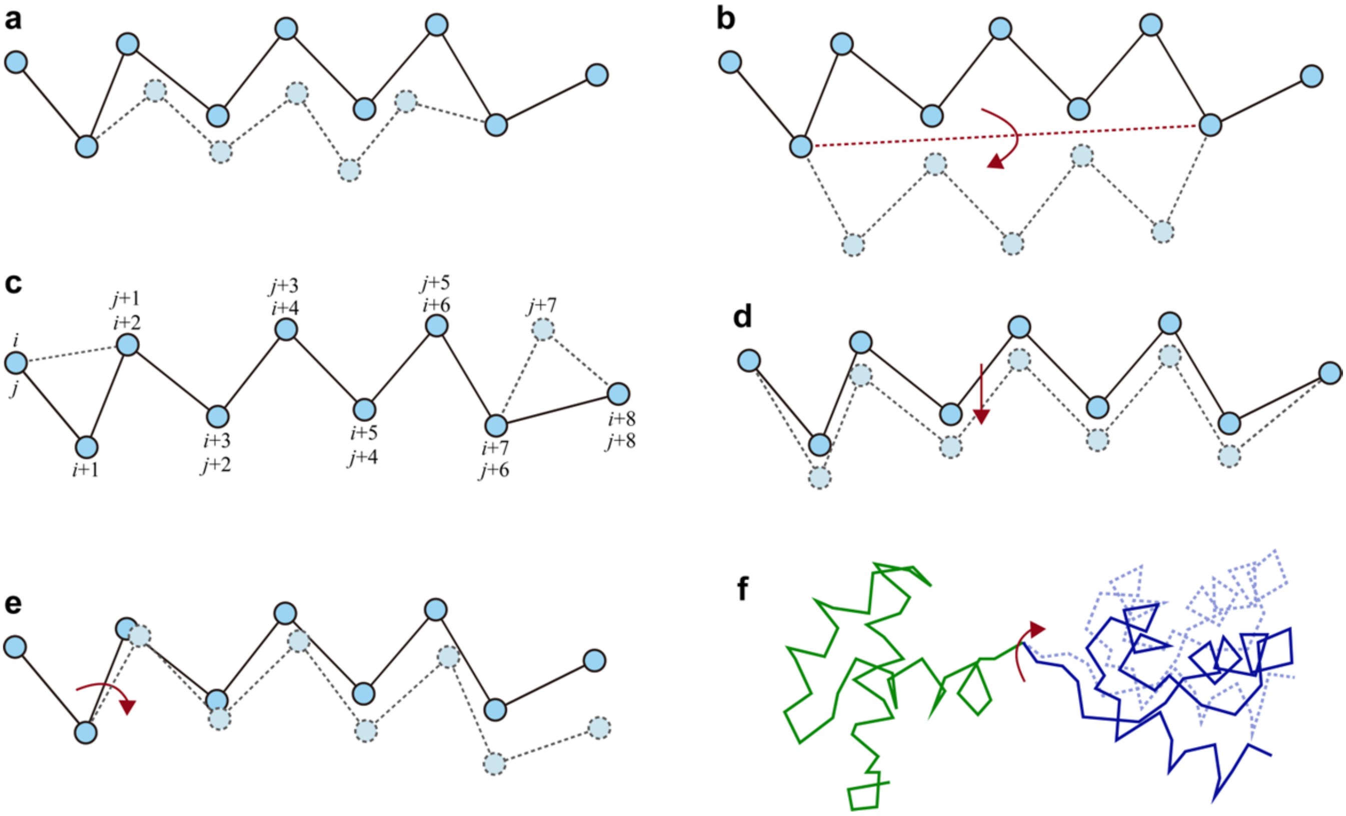


**Supplementary Figure 14**

Movements for the flexible domain assembly and refinement.

(**a**) LMProt[6] perturbation. (**b**) Segment rotation around the axis connecting two termini of the segment. (**c**) Conformational shift of segments along the sequence. (**d**) Rigid-body segment translation. (**e**) rigid-body tail rotation. (**f**) Rigid-body domain-level translation and rotation.

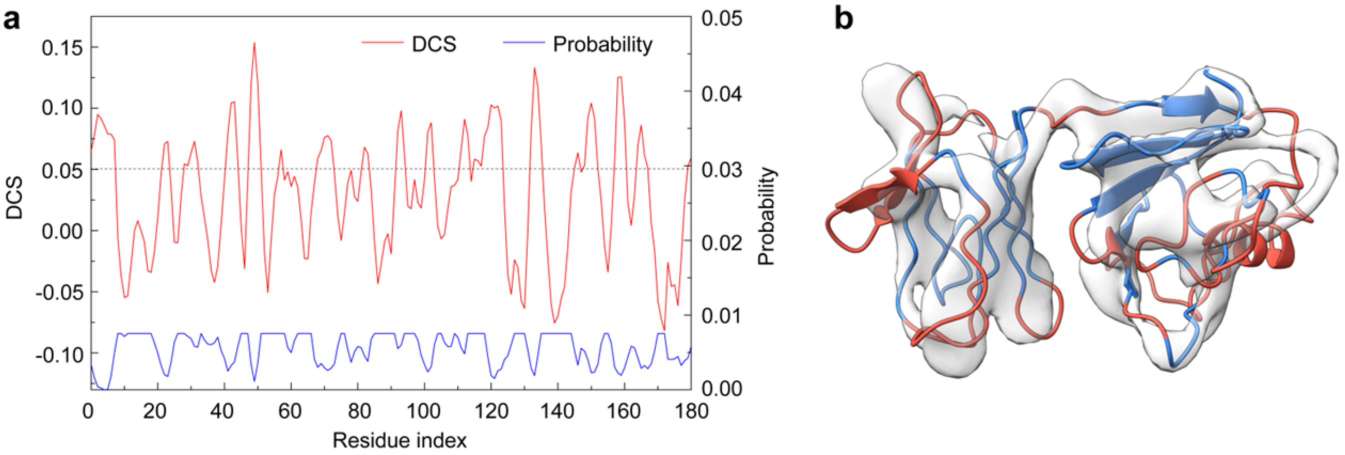


**Supplementary Figure 15**

Example of regions determination for remodeling and refinement.

(**a**)The local density correlation score of each residue in an example protein (PDB 1wv3A) and the corresponding probability of each residue to be selected for remodeling. (**b**) The 3D structure of 1wv3A superposed into the density map, where red regions are residues with local density score <0.05.

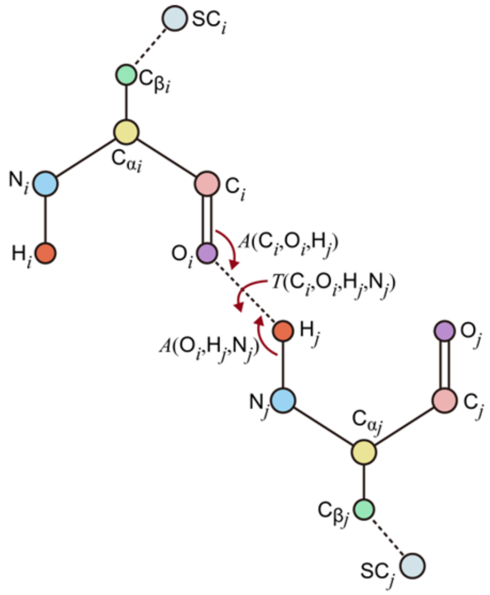


**Supplementary Figure 16**

Definition of H-O distance, inner angles, and torsion angle for hydrogen-bond.

### Supplementary Tables

**Supplementary Table 1.** Summary of final models constructed from experimental domain structures on 357 test proteins using simulated density maps. Bold font highlights the best results from each category.

| | Method | TM-score | RMSD (Å) |
|---|---|---|---|
| 2dom[a] (*N*=166) | MDFF | 0.89 | 4.7 |
| | Rosetta | 0.86 | 4.8 |
| | DEMO-EM | **0.99** | **0.5** |
| 2dis[b] (*N*=81) | MDFF | 0.87 | 5.1 |
| | Rosetta | 0.85 | 5.4 |
| | DEMO-EM | **0.99** | **0.6** |
| 3dom[c] (*N*=69) | MDFF | 0.76 | 10.8 |
| | Rosetta | 0.72 | 11.0 |
| | DEMO-EM | **0.99** | **0.7** |
| m4dom[d] (*N*=41) | MDFF | 0.63 | 20.2 |
| | Rosetta | 0.55 | 21.9 |
| | DEMO-EM | **0.99** | **0.8** |
| All (*N*=357) | MDFF | 0.83 | 7.7 |
| | Rosetta | 0.79 | 8.1 |
| | DEMO-EM | **0.99** | **0.6** |

[a] protein with 2 domains.
[b] protein with discontinuous domain which contains 2 or more segments from separate regions of the query sequence.
[c] protein with 3 domains.
[d] protein with 4 or more domains.

**Supplementary Table 2.** Summary of final models constructed from I-TASSER domain models using simulated density maps. Bold font highlights the best results from each category.

| | Method | TM-score | RMSD(Å) | TM-score (domain)[a] | RMSD(Å) (domain)[b] |
|---|---|---|---|---|---|
| 2dom (*N*=116) | MDFF | 0.61 | 13.9 | 0.66 | 5.7 |
| | Rosetta | 0.61 | 13.7 | 0.62 | 6.8 |
| | DEMO-EM | **0.87** | **4.8** | **0.83** | **3.8** |
| 2dis (*N*=41) | MDFF | 0.58 | 15.7 | 0.66 | 7.7 |
| | Rosetta | 0.56 | 15.9 | 0.60 | 8.7 |
| | DEMO-EM | **0.83** | **8.0** | **0.80** | **6.1** |
| 3dom (*N*=47) | MDFF | 0.48 | 19.0 | 0.66 | 8.2 |
| | Rosetta | 0.43 | 19.7 | 0.51 | 8.6 |
| | DEMO-EM | **0.85** | **5.8** | **0.83** | **3.2** |
| m4dom (*N*=25) | MDFF | 0.36 | 32.0 | 0.62 | 8.2 |
| | Rosetta | 0.27 | 34.2 | 0.38 | 12.0 |
| | DEMO-EM | **0.82** | **8.5** | **0.80** | **4.1** |
| All (*N*=229) | MDFF | 0.55 | 17.2 | 0.65 | 6.2 |
| | Rosetta | 0.53 | 17.5 | 0.57 | 8.1 |
| | DEMO-EM | **0.85** | **6.0** | **0.83** | **4.1** |

[a] TM-score of individual domain models in full-length models.
[b] RMSD of individual domains models in full-length models.

**Supplementary Table 3.** Summary of refinement results for the full length models generated by the rigid-body assembly of DEMO-EM using I-TASSER predicted domain structures. Bold font highlights the best results.

| Method | TM-score | RMSD(Å) | TM-score (domain) | RMSD(Å) (domain) |
|---|---|---|---|---|
| Initial Model[a] | 0.78 | 8.4 | 0.76 | 5.0 |
| MDFF-DEMO-EM[b] | 0.82 | 7.1 | 0.80 | 4.5 |
| Rosetta-DEMO-EM[c] | 0.81 | 7.3 | 0.77 | 4.6 |
| DEMO-EM | **0.85** | **6.0** | **0.83** | **4.1** |

[a] Full length models generated by the rigid-body assembly of DEMO-EM.
[b] MDFF refinement for the full length models by the rigid-body assembly of DEMO-EM.
[c] Rosetta refinement for the full length models by the rigid-body assembly of DEMO-EM.

**Supplementary Table 4.** Domain definitions of the 6 proteins in SARS-CoV-2 genome by FUpred[7]. Different domains are separated by semicolons, and different segments of the discontinuous domain are separated by commas

| EMD code | Resolution(Å) | PBD code | Domain definition |
|---|---|---|---|
| EMD-21375 | 3.46 | 6vsbC | 1-290;291-320,591-700;321-327,529-590;328-528; 701-717, 1072-1146; 718-1071; |
| EMD-11007 | 2.9 | 6yytB | 1-125;126-198; |
| EMD-22160 | 3.5 | 6xezE | 1-100;101-235;236-439;440-605; |
| EMD-22136 | 2.9 | 6xdcA | 1-141;142-284; |
| EMD-11007 | 2.9 | 6yytC | 1-83; |
| EMD-11007 | 2.9 | 6yytA | 1-265;266-388;389-407,445-812;408-444,813-932; |

**Supplementary Table 5.** Statistic values to determine coordinates of O, $C_\beta$, H, and side-chain center of mass (SC) according to their relative positions to the three backbone atoms (N, $C_\alpha$, and C), where *D*, *T*, and *A* indicate the distance, torsion angle, and inner angle, respectively

| | | | | | |
|---|---|---|---|---|---|
| $O_i$ $i \in [1, L-1]$ | $D(O_i,C_i)$ (Å) | 1.229 | $H_i$ $i \in [2, L]$ | $D(N_i,H_i)$ (Å) | 0.987 |
| | $T(C_{\alpha_i},C_i,O_i,N_{i+1})$ (°) | 179.672 | | $T(C_{i-1},N_i,C_{\alpha_i},H_i)$ (°) | 179.817 |
| | $A(C_{\alpha_i},C_i,O_i)$ (°) | 120.098 | | $A(H_i,N_i,C_{\alpha_i})$ (°) | 119.255 |
| $O_i$ $i = L$ | $D(O_i,C_i)$ (Å) | 1.244 | $H_i$ $i = 1$ | $D(N_i,H_i)$ (Å) | 0.987 |
| | $T(C_{\alpha_i},C_i,O_i,N_i)$ (°) | 0 | | $T(C_i,N_i,C_{\alpha_i},H_i)$ (°) | 60 |
| | $A(C_{\alpha_i},C_i,O_i)$ (°) | 119.494 | | $A(H_i,N_i,C_{\alpha_i})$ (°) | 116.345 |
| $C_\beta$ | Residue type specific: https://zhanglab.ccmb.med.umich.edu/DEMO-EM/potential/CB_position.txt | | | | |
| SC | Residue type specific: https://zhanglab.ccmb.med.umich.edu/DEMO-EM/potential/SG_position.txt | | | | |

**Supplementary Table 6.** Van der Waals radius parameters from CHARMM[8] are used to count for excluded volume interaction.

| | $C_\alpha$ | N | C | O | $C_\beta$ | SC |
|---|---|---|---|---|---|---|
| $C_\alpha$ | 3.6 | 2.3 | 3.7 | 2.9 | 3.5 | 1.0 |
| N | 3.7 | 2.5 | 3.5 | 2.5 | 3.5 | 1.0 |
| C | 2.3 | 1.2 | 2.7 | 2.7 | 2.3 | 1.0 |
| O | 2.5 | 2.1 | 2.4 | 2.3 | 2.6 | 1.0 |
| $C_\beta$ | 3.5 | 2.3 | 3.5 | 2.8 | 3.3 | 1.0 |
| SC | 1.0 | 1.0 | 1.0 | 1.0 | 1.0 | 1.0 |

**Supplementary Table 7.** Mean and standard deviation of four hydrogen-bond features in α-helix and β-sheet structures.

| | $D(O_i, H_j)$ (Å) | $A(C_i, O_i, H_j)$ (°) | $A(O_i, H_j, N_j)$ (°) | $T(C_i, O_i, H_j, N_j)$ (°) |
|---|---|---|---|---|
| Helix, *j*=*i*+4 | 2.00/0.53 | 147/10.58 | 159/11.25 | 160/25.36 |
| Helix, *j*=*i*+3 | 2.85/0.32 | 89/7.70 | 111/8.98 | -160/7.93 |
| Parallel | 2.00/0.30 | 155/11.77 | 164/11.29 | 180/68.96 |
| Antiparallel | 2.00/0.26 | 151/12.38 | 163/11.02 | -168/69.17 |

### Supplementary Texts

**Supplementary Text 1. Implementation of MDFF and Rosetta programs**

MDFF is cryo-EM density-map guided protein structure fitting and refinement program though a combined search process of Monte Carlo simulation, conjugate-gradients minimization, and simulated annealing molecular dynamics simulations[3]. Starting from the initial full-length model built by matching each domain model into density maps using Situs[5], MDFF was carried out by a rigid-body assembly with domain restraints for 70ps, a flexible fitting for 1,000ps, and a final energy minimization step. Following the protocol described previously[9], the electron density term with a weight of 1, 0.3, and 10 was used in three steps for the flexible fitting.

Rosetta builds cryo-EM models also through Monte Carlo simulations as guided by the Rosetta all-atom force field[4]. Started with the same initial models constructed from Situs[5], Rosetta was performed according to the protocol of cyro-EM density based model refinement using iterative local rebuilding[4], which is described in https://faculty.washington.edu/dimaio/files/rosetta_density_tutorial_aug18_2.pdf.
